# Airway Secretory Cells Contain Both a Perinuclear Golgi Ribbon and Dispersed Golgi Satellites

**DOI:** 10.1101/2025.04.11.648270

**Authors:** Oanh N. Hoang, Colin E. Chan, Joshua M. Brenner, Denisse Leza-Rincon, Ana M. Jaramillo, Brendan Dolan, Adam W. Aziz, Rodolfo A. Cardenas, Gerardo J. Cardenas, Eduardo D. Galvez, Reid T. Powell, Leoncio Vergara, Harry Karmouty-Quintana, Jesper M. Magnusson, Gunnar C. Hansson, Roberto Adachi, John D. Dickinson, Christopher M. Evans, Justin A. Courson, Alan R. Burns, Michael J. Tuvim, Burton F. Dickey

## Abstract

**Rationale:** Finely tuned production and secretion of the polymeric mucins MUC5AC and MUCB are required for lung health, but knowledge of many details between their translation and their packaging into secretory granules is lacking.

**Objectives:** To analyze the structure and function of the Golgi apparatus, a key site of mucin glycosylation, folding, polymerization and packaging, in airway epithelial secretory cells.

**Methods:** Lung tissue was obtained from mice stimulated or not with IL-13 to upregulate mucin production, and from normal human lungs. Golgi elements in mouse and human tissue were imaged by high-resolution immunofluorescence microscopy and electron microscopy. Tissue from mice with deletion of both polymeric mucins was also examined.

**Measurements and Main Results:** By immunofluorescence microscopy, both mouse and human airway secretory cells contained ∼100 dispersed puncta labeled by markers of medial and trans Golgi cisternae and the trans-Golgi network (TGN), but only a few perinuclear puncta were labeled by markers of cis-Golgi cisternae. By electron microscopy, secretory cells contained both a perinuclear Golgi ribbon and numerous dispersed Golgi stacks, termed satellites. In mucous metaplastic cells, satellites were concentrated among immature mucin granules. Increasing mucin production by cytokine stimulation did not increase the number of TGN puncta, nor did preventing polymeric mucin production by gene deletion reduce TGN puncta.

**Conclusions:** Mucin-producing airway secretory cells express an unusual Golgi structure consisting of a conventional perinuclear Golgi ribbon as well as dispersed satellites. While the Golgi satellites are likely an adaptation for mucin production and packaging, their presence is specified developmentally, independent of mucin production.

## INTRODUCTION

The physical properties of airway mucus arise from interactions in the airway lumen of the secreted polymeric mucins, MUC5AC and MUC5B, with water and salts (1–3).

These mucins are enormous glycoproteins, with sizes as monomers greater than two million Daltons, and they polymerize into linear chains containing tens of monomers. The huge size of mucin polymers is required for effective mucociliary clearance (4), but it places great demands on airway secretory cells that leads to proteostasis stress (5). Finely tuned production and secretion of MUC5AC and MUC5B are required for lung health. The absence of MUC5AC in mice results in impaired trapping of helminthic larvae migrating through the lungs (6), but hyperexpression of MUC5AC combined with rapid secretion in mice and humans causes mucus plugging of airways (7–11). The absence of MUC5B in mice and in humans results in impaired mucociliary clearance leading to airway inflammation, infection, and injury (12, 13). However, hyperexpression of MUC5B in mice increases susceptibility to bleomycin-induced lung injury (14, 15), and hyperexpression in humans is an important contributor to fibrotic interstitial lung diseases (16–19).

In view of the clinical importance of airway mucin production and secretion, the transcriptional control of MUC5AC and MUC5B synthesis and the mechanism of exocytic mucin secretion have been analyzed in detail (20–22). However, intermediate steps in mucin production, such as post-translational modifications including N-terminal polymerization and late stages of glycosylation, trafficking from endoplasmic reticulum through the Golgi apparatus, and packaging into secretory granules, have received less attention. Recently, we showed that most secretory granules contain both MUC5AC and MUC5B tightly interdigitating (23), similar to the packaging of two different mucins within single granules in *Drosophila* salivary glands (24). As our next goal in analyzing intermediate steps in mucin production, we sought to examine the packaging of mucins within granules together with exocytic proteins on the granule surface, such as VAMP-8 and Synaptotagmin-2 (25, 26), that make the granule competent for secretion. Co-packaging of mucins and exocytic proteins could be expected to occur in the trans-Golgi network (TGN), where assembly of secretory granules has been studied in multiple endocrine and exocrine cells (27, 28). However, when we visualized the TGN of airway secretory cells by immunofluorescence microscopy, we observed numerous widely dispersed puncta rather than concentrated perinuclear immunofluorescence as would be expected for a classical Golgi ribbon.

A Golgi apparatus is found in essentially all eukaryotic cells, but its appearance is highly variable across phyla. In most vertebrate cells, the Golgi apparatus has a stereotypic structure consisting of several cisternal stacks linked laterally into a single ribbon (29).

In polarized epithelial cells, the Golgi ribbon is usually located just apical to the nucleus. While a ribbon is the most common Golgi structure in vertebrate cells, other Golgi structures are sometimes observed in highly specialized cells. For example, Golgi mini-stacks termed satellites are seen in neuronal dendrites where they are thought to enable local protein synthesis without requiring lengthy transport from the cellular soma (30–32). Another example is the occurrence of dispersed mini-stacks within endothelial cells where they are thought to enable the assembly of very large von Willibrand factor polymers into secretory Weibel-Palade bodies (33–35). Besides its key roles in protein synthesis and trafficking, the Golgi apparatus is increasingly recognized for providing a platform for multiple cellular processes including cytoskeletal organization, sensing of proteostasis stress, metabolism, autophagy, inflammation, and apoptosis (29, 31).

These activities can be modulated by changes in Golgi structure, so knowledge of the structure and function of the Golgi apparatus in lung parenchymal cells is important to understand lung health and pathophysiology. Here we determine the ultrastructure and distribution of Golgi stacks in airway epithelial secretory cells, their association with polymeric secretory mucins, and their developmental specification.

## METHODS

Briefly, wild-type C57BL/6 mice, challenged or not with cytokines to induce mucous metaplasia, were studied at MD Anderson Cancer Center under approved institutional protocols. MUC5AC/MUC5B double deletant mice were generated at the University of Colorado under approved institutional protocols. Normal human tissue was obtained at the University of Texas Health Science Center at Houston and the University of Gothenburg under approved institutional protocols. For microscopic analysis, widefield deconvolution immunofluorescence microscopy was performed at MD Anderson Cancer Center, electron microscopy at the University of Houston College of Optometry, and high resolution Airyscan immunofluorescence microscopy at the University of Gothenburg. Detailed methods are provided in the Online Data Supplement.

## RESULTS

### The trans-Golgi network (TGN) of mouse airway secretory cells is widely dispersed

As an initial step towards visualizing the assembly of mucin secretory granules, we imaged the TGN in airway epithelium by brightfield fluorescence microscopy using an antibody against TGN46. Rather than finding a few fluorescent puncta close near the nucleus as expected for the Golgi ribbon of a typical mammalian cell (32), we observed numerous puncta widely distributed throughout the cytoplasm of airway secretory cells (not shown). In a pilot study to confirm the large number of TGN elements and their widespread distribution, we performed laser confocal immunofluorescence microscopy of naïve, uninflamed airway epithelial cells and airway cells with mucous metaplasia due to IL-13 instillation (Figure E1). Using a volumetric algorithm to estimate the number of TGN elements per cell (see Methods), we found a mean of 87 puncta in naïve secretory cells compared to 11 in ciliated cells, and 113 puncta in metaplastic secretory cells compared to 11 in ciliated cells. Besides the difference in the number of puncta, TGN46 was distributed throughout the cytoplasm of secretory but not ciliated cells, extending close to the apical surface of tall metaplastic secretory cells distended with mucin granules (Figure E1B).

To further assess these initial findings, we imaged mouse axial bronchi by widefield deconvolution immunofluorescence microscopy using a similar volumetric algorithm.

This showed a mean of 144 TGN46 puncta in naïve secretory cells compared to 45 in ciliated cells, and 206 puncta in metaplastic secretory cells compared to 72 in ciliated cells (Figure 1). Again, TGN46 was distributed throughout the cytoplasm of secretory but not ciliated cells, extending to the apical pole of tall metaplastic secretory cells (Figures 1 and E2). For comparison to a cell with a more conventional Golgi structure, we imaged submucosal fibroblasts marked by the expression of peptidase inhibitor 16 (PI16, Figure 1D). The fibroblasts showed TGN46 puncta adjacent to one side of the nucleus interspersed with GM130 puncta that mark the cis-Golgi, consistent with a Golgi ribbon containing complete stacks spanning from cis to trans cisternae. Thus, mouse airway secretory cells show a larger number and wider dispersion of TGN elements than adjacent ciliated epithelial cells and submucosal fibroblasts.

**Figure 1.**
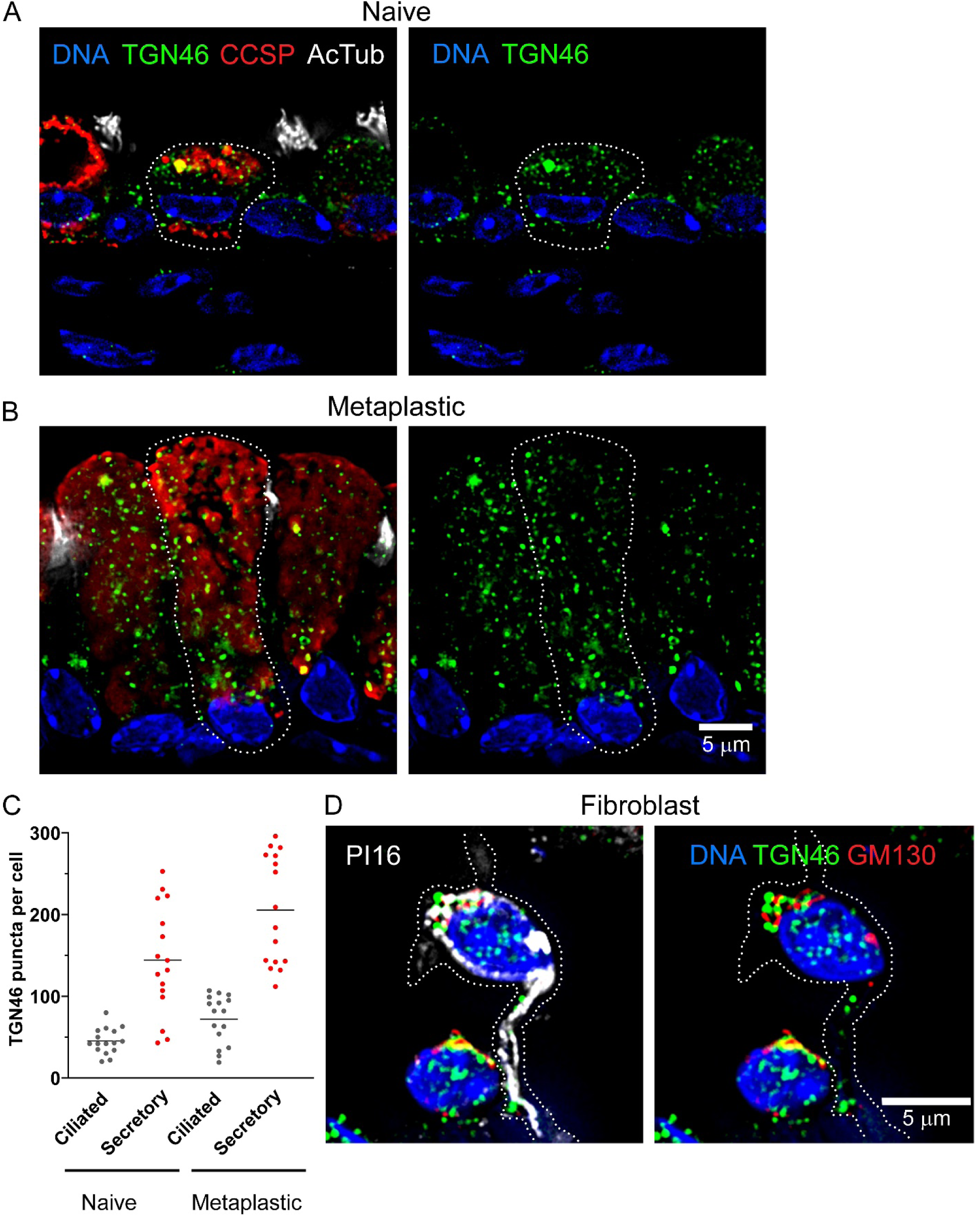
Numerous widely distributed trans-Golgi network (TGN) elements in mouse airway secretory cells. (*A)* Immunofluorescence deconvolution microscopy using antibodies against CCSP to mark secretory cells, acetylated tubulin (AcTub) to mark ciliated cells, TGN46 to mark the trans-Golgi network, and DAPI to stain nuclei in the axial bronchi of mice not exposed (naïve) to cytokines. The scale bar in *B* also applies here. (*B*) Microscopy as in *A*, but in mice with mucous metaplasia due to treatment with IL-13. (*C*) TGN puncta were enumerated in 16 cells from 3 mice in each group. Selected comparisons were subjected to Mann-Whitney test with Bonferroni correction, which set significance at *p* < 0.0125. For naïve ciliated vs naïve secretory, *p* < 0.0001; for metaplastic ciliated vs metaplastic secretory, *p* < 0.0001; for naïve ciliated vs metaplastic ciliated, *p* = 0.01; for naïve secretory vs metaplastic secretory, *p* = 0.02. (D) Microscopy as in *A* of a submucosal fibroblast marked by the expression of PI16 (left panel), and showing TGN46 puncta aggregated with puncta of the cis-Golgi marker GM130 adjacent to a single pole of the nucleus (right panel).

### Mouse airway secretory cells contain both a conventional Golgi ribbon and dispersed Golgi satellites by electron microscopy (EM)

To characterize the structure of the Golgi apparatus in mouse airway secretory cells at higher resolution, we used transmission EM. Golgi stacks linked laterally in ribbons that are conventional for vertebrate cells were observed in both naïve and mucous metaplastic airway secretory cells (Figure 2A-B). Most often these ribbons were located close to nuclei on the apical side. In addition to ribbons, isolated Golgi stacks were observed throughout the cytoplasm, most commonly apical or lateral to the nucleus rather than basal. These scattered Golgi stacks are termed satellites in reference to Golgi stacks that are found distal from Golgi ribbons in other cell types such as neurons (32). In naïve secretory cells (Figure 2A), the Golgi satellites were observed throughout the apical half of cells that contained very few mucin granules (note, atypical round mitochondria with minimal internal structure should not be mistaken for secretory granules, see Figures E3 and E4 and Discussion). In metaplastic secretory cells (Figure 2B), the Golgi satellites were mostly observed among densely packed immature mucin granules in the middle third of cells; few satellites were observed among the mature mucin granules in the apical third of cells or in the basal third that included the nucleus. Enlarged and annotated images of the naïve and metaplastic cells in Figure 2A-B are provided in the Online Supplement along with a gallery of additional EM images (Figures E3 and E4) and images from serial block-face scanning EM of several metaplastic secretory cells (Figure E5, Videos 1 and 2).

**Figure 2.**
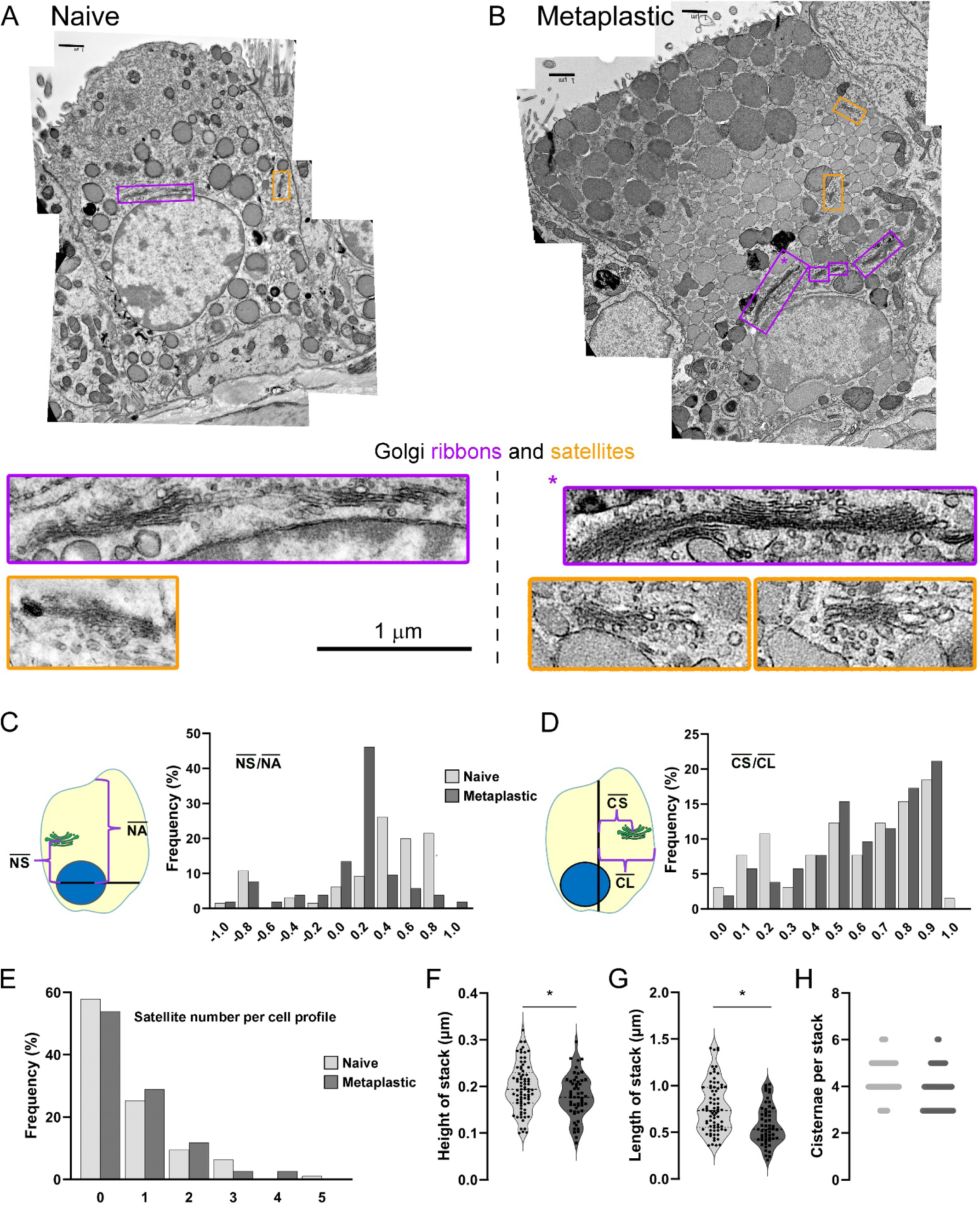
Electron microscopic analysis of Golgi structure in mouse airway secretory cells. Cellular profiles were compiled from multiple electron micrographs of secretory cells in mouse axial bronchi to include apical, basal, and lateral plasma membranes as well as a portion of the nucleus. Cellular profiles were compiled for 95 naïve (uninflamed) cells and 76 metaplastic cells. *(A)* Representative airway secretory cell of a naïve mouse. The Golgi ribbon is outlined in magenta, a Golgi satellite is outlined in orange, and both are also shown at higher magnification. All scale bars in *A* and *B* are 1 µm. *(B)* Representative metaplastic secretory cell in the airway of a mouse treated with IL-13. The Golgi ribbon and satellites are outlined as in *A*, but only the portion of the ribbon indicated by an asterisk is shown at higher magnification*. (C)* Frequency distribution of the vertical position of satellites relative to the nucleus (NS) as a fraction of the distance from the center of the nucleus to the middle of the apical membrane (NA). Intervals extend 0.1 units more and less than the listed fractions, and positions below the center of the nucleus are negative. The vertical distribution of satellites differs between naïve and metaplastic cells, *p*<0.0001. *(D)* Frequency distribution of the lateral position of satellites relative to the center of the cell (CS) as a fraction of the distance from the center of the cell to the lateral membrane on that side at the level of the satellite (CL). Intervals extend 0.05 units more and less than the listed fractions. The horizontal distribution of satellites does not differ between naïve and metaplastic cells, *p* = 0.707. *(E)* Frequency distribution of the number of satellites per cellular profile shows no difference between naïve and metaplastic cells, *p* > 0.99. The morphology of satellites in naïve and metaplastic secretory cells was compared based on height *(F; p=*0.046), length *(G; p* < 0.01), and number of cisternae per satellite*(H; p*<0.01). Statistical comparisons were done by the Kolmogorv-Smirnov test (*C-E*) and Mann-Whitney *U* test (*F-H*).

### Quantitative ultrastructural analysis of mouse Golgi satellites

Our qualitative observations were extended quantitatively by measuring the distribution of satellites relative to the apical and lateral cell borders, the number of satellites, and the dimensions of satellites in EM cellular profiles of 95 naïve cells and 76 metaplastic cells. Criteria for the identification of secretory cells and the compilation of EM images to generate complete cross-sectional profiles are provided in the Online Supplemental Methods. In naïve cells, satellites were found mostly at a fractional distance from the apical membrane of 0.4-0.8, whereas in metaplastic cells, satellites were found mostly at a fractional distance of 0-0.4 (Figure 2C). This analysis is consistent with the qualitative observation of the redistribution of satellites from the apical region in naïve cells to the middle region in metaplastic cells. There was a gradient of slightly increasing satellite number towards the lateral borders of both naïve and metaplastic cells with no observable redistribution with metaplasia (Figure 2D).

The number of satellites visible in an EM cellular profile varied from 0 to 5, with no significant difference in the means of 0.68 in naïve cells and 0.71 in metaplastic cells (Figure 2E). Based upon our estimate of the volume fraction of an airway cell sampled by an EM cellular profile, we calculate a mean total satellite number of 55.5 per naïve cell and 57.9 per metaplastic cell (see the Results section of the Online Supplement). The mean height of satellites in naïve cells was 0.20 µm and in metaplastic cells was 0.18 µm (Figure 2F), the mean length of satellites in naïve cells was 0.78 µm and in metaplastic cells was 0.58 µm (Figure 2G), and the mean number of cisternae in satellites in naïve cells was 4.4 and in metaplastic cells was 3.9 (Figure 2H). Thus, while there is a clear redistribution of satellites during mucous metaplasia towards the middle third of secretory cells where numerous immature mucin granules are concentrated, there was no significant difference between naïve and metaplastic cells in the number of satellites and only small differences in their dimensions.

### Cis-Golgi cisternae are concentrated in the ribbon whereas trans-Golgi cisternae are present in both the ribbon and satellites in mice

To determine whether Golgi stacks in the ribbon and satellites had the same or a different composition, we performed immunofluorescence deconvolution microscopy using antibodies against the cis-Golgi marker GM130 and the trans-Golgi marker GRASP55. GM130 was concentrated in the perinuclear region of both naïve and metaplastic secretory cells, similar to its distribution in neighboring ciliated cells and consistent with predominant localization in the Golgi ribbon (Figures 3 and E6). In contrast, GRASP55 was observed throughout the cytoplasm of secretory cells, consistent with localization in both the ribbon and satellites. GM130 and GRASP55 were observed to colocalize in the perinuclear region, consistent with their presence within a single Golgi stack in ribbons (yellow color in Figures 3 and E6). To observe the localization of GM130 and TGN46 in relation to mucin granules, antibodies to MUC5B and MUC5AC were used. GM130 was seen in a perinuclear location not overlapping MUC5B in either naïve or metaplastic cells (Figure E7A-B). In contrast, TGN46 was seen throughout the cytoplasm, including in the region of MUC5B granules in naïve and metaplastic cells (Figure E7C-D), and the region of MUC5AC in metaplastic cells (Figure E7F).

**Figure 3.**
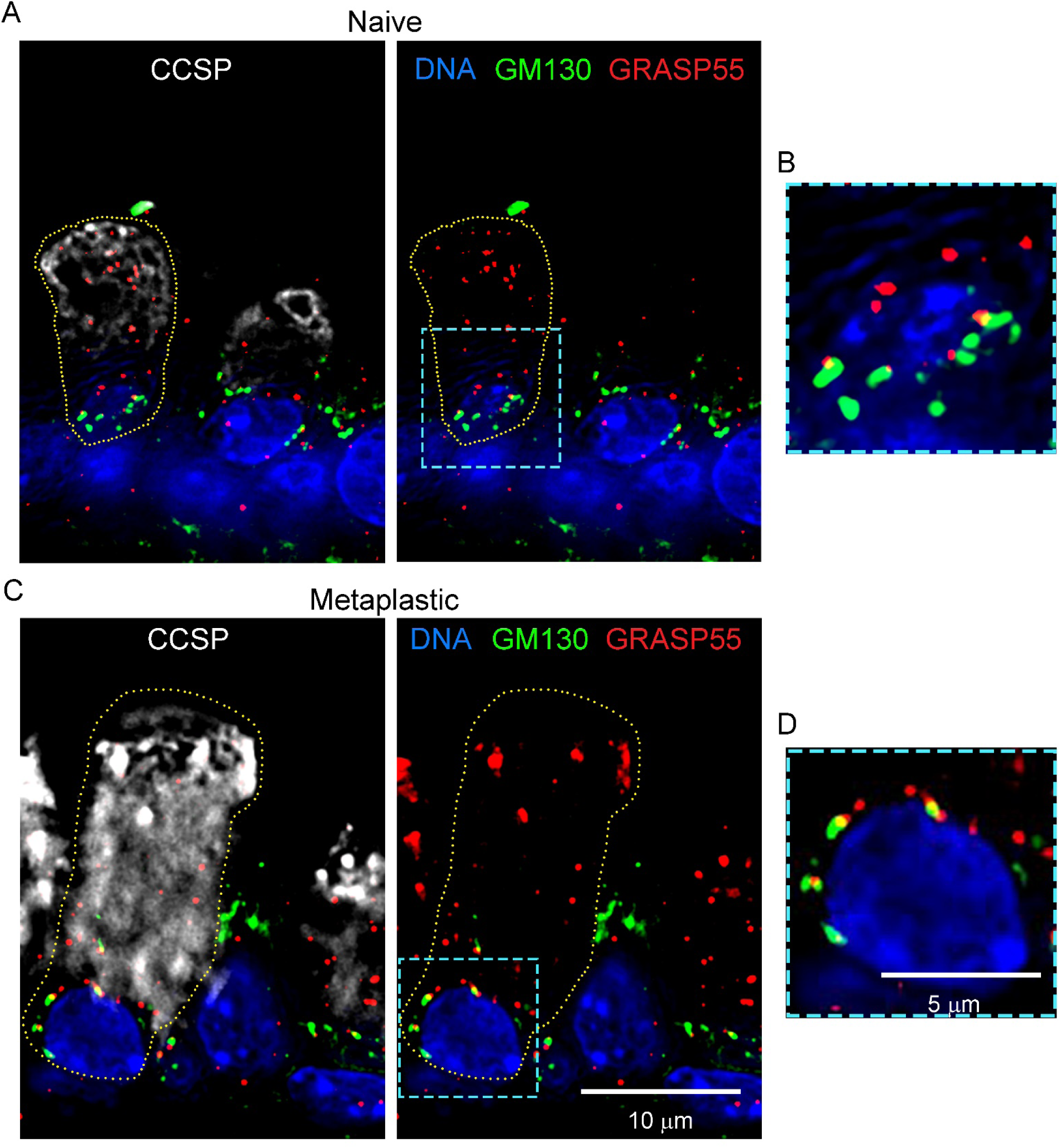
Localization of cis-Golgi and trans-Golgi cisternae in mouse airway secretory cells. *(A)* Immunofluorescence deconvolution microscopy using antibodies against CCSP to mark secretory cells, GM130 to mark cis-Golgi cisternae, GRASP55 to mark trans-Golgi cisternae, and DAPI stain to mark nuclei in the axial bronchus of naïve mice. *(B)* The perinuclear area outlined in *A* is shown at higher magnification. *(C)* Microscopy as in *A*, but in mice with mucous metaplasia due to treatment with IL-13. *(D)* The perinuclear area outlined in *C* is shown at higher magnification.

### TGN and trans-Golgi cisternae in human airway secretory cells are widely dispersed

To determine whether the structure of the Golgi apparatus in human airway secretory cells is similar to that in mice, we probed sections of proximal and distal human airways with antibodies against TGN46, GRASP55, and GM130 using deconvolution immunofluorescence microscopy. In proximal airways, antibodies against TGN46 showed widely dispersed TGN from the nucleus all the way to the apical plasma membrane of tall secretory cells (Figure 4A). Neighboring ciliated cells did not show similarly dispersed TGN46 staining. In distal airways, epithelial cells were considerably shorter, but TGN46 staining again extended apically from the nucleus (Figure 4B). In both proximal and distal airway secretory cells, TGN46 morphology was both punctate and tubular, with many tubules apparently wrapped around CCSP-containing secretory granules (Figure 4B inset) and around mucin granules containing MUC5AC or MUC5B or both mucins (Figure E8). In contrast to the dispersed expression of TGN46 in airway secretory cells, submucosal fibroblasts identified by MEOX2 staining showed TGN46 in just one or two perinuclear puncta associated with the cis-Golgi marker GM130, consistent with their exclusive localization to a Golgi ribbon (Figure 4C).

**Figure 4.**
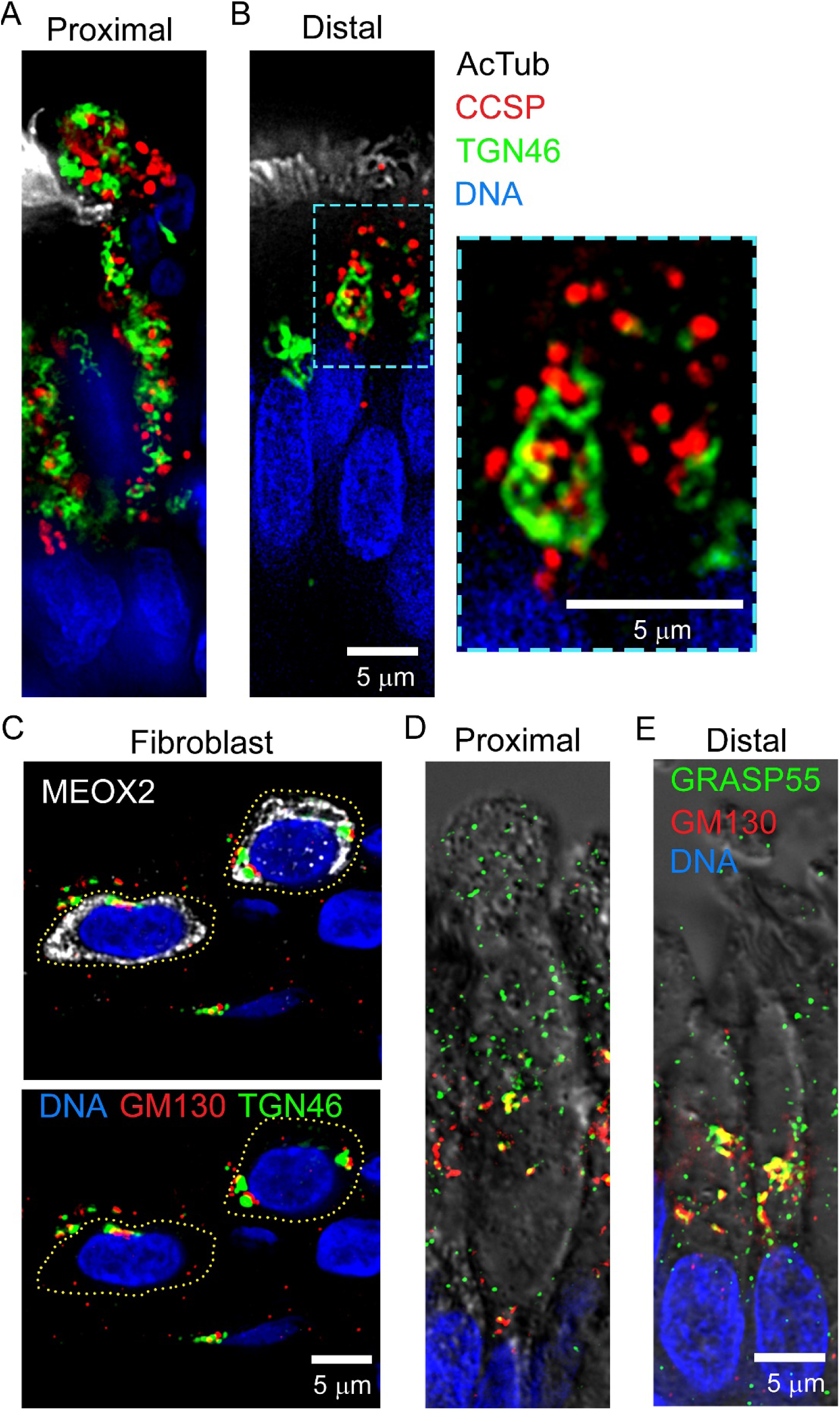
Localization of Golgi cisternae in human airway secretory cells. *(A)* Immunofluorescence deconvolution microscopy using antibodies against acetylated tubulin (AcTub) to mark ciliated cells, CCSP to mark secretory cells, TGN46 to mark the trans-Golgi network, and DAPI to mark nuclei in a proximal human airway. *(B)* Microscopy as in *A*, but in a distal human airway. The area marked in the secretory cell is shown at higher magnification on the right. *(C)* Microscopy as in *A*, but using antibodies against MEOX2 to mark fibroblasts and antibodies against GM130 to mark cis-Golgi cisternae. The MEOX2 channel is left out of the bottom image to better visualize GM130 and TGN46. *(D-E)* Microscopy as in *A-B*, but using antibodies against GRASP55 to mark trans-Golgi cisternae and against GM130 to mark cis-Golgi cisternae. Fluorescence images are superimposed on differential interference contrast images.

To determine whether there is a spatial dissociation between cis and trans Golgi elements in human airway secretory cells as in mice, we used antibodies against GRASP55 that localizes to the trans-Golgi and against GM130 that localizes to the cis-Golgi. In proximal airway secretory cells, GRASP55 was widely dispersed throughout the cytoplasm, similar to the distribution of TGN46 (Figure 4D). In the shorter distal airway secretory cells, GRASP55 was also observed in both dispersed and perinuclear locations, though the dispersed element was less abundant than in proximal cells (Figure 4E). In contrast to GRASP55, GM130 was concentrated near the nucleus in both proximal and distal airways (Figure 4D-E), often colocalized with GRASP55 consistent with their joint presence in a Golgi ribbon (yellow color). Thus, similar to mice, human airway secretory cells show TGN and trans-Golgi elements to be widely dispersed, consistent with their presence in both satellites and the ribbon, whereas cis-Golgi elements are concentrated near the nucleus, consistent with their presence more exclusively in the ribbon.

### Human airway secretory cells contain dispersed Golgi satellites by EM

To confirm that dispersed trans-Golgi elements observed by immunofluorescence microscopy of human airway secretory cells represent Golgi mini-stacks similar to those in mice, we performed transmission EM of human airway tissue and cultured human airway epithelial cells. In tissue sections, proximal airway secretory cells showed Golgi mini-stacks in close apposition to mucin granules in the middle third of cells (Figure 5A). Zinc iodide – osmium tetroxide (ZIO) fixation/staining confirmed the abundance of dispersed Golgi elements in mucin-containing secretory cells (Figure 5B). In cultured human airway epithelial cells, dispersed Golgi elements were observed both in unstained (Figure 5C) and ZIO-stained (Figure 5D) specimens, and perinuclear Golgi ribbons were also seen (Figure 5D). Golgi mini-stacks were also observed in close apposition to mucin granules in stained specimens of submucosal gland mucous cells (Figures 5E-F and E9).

**Figure 5.**
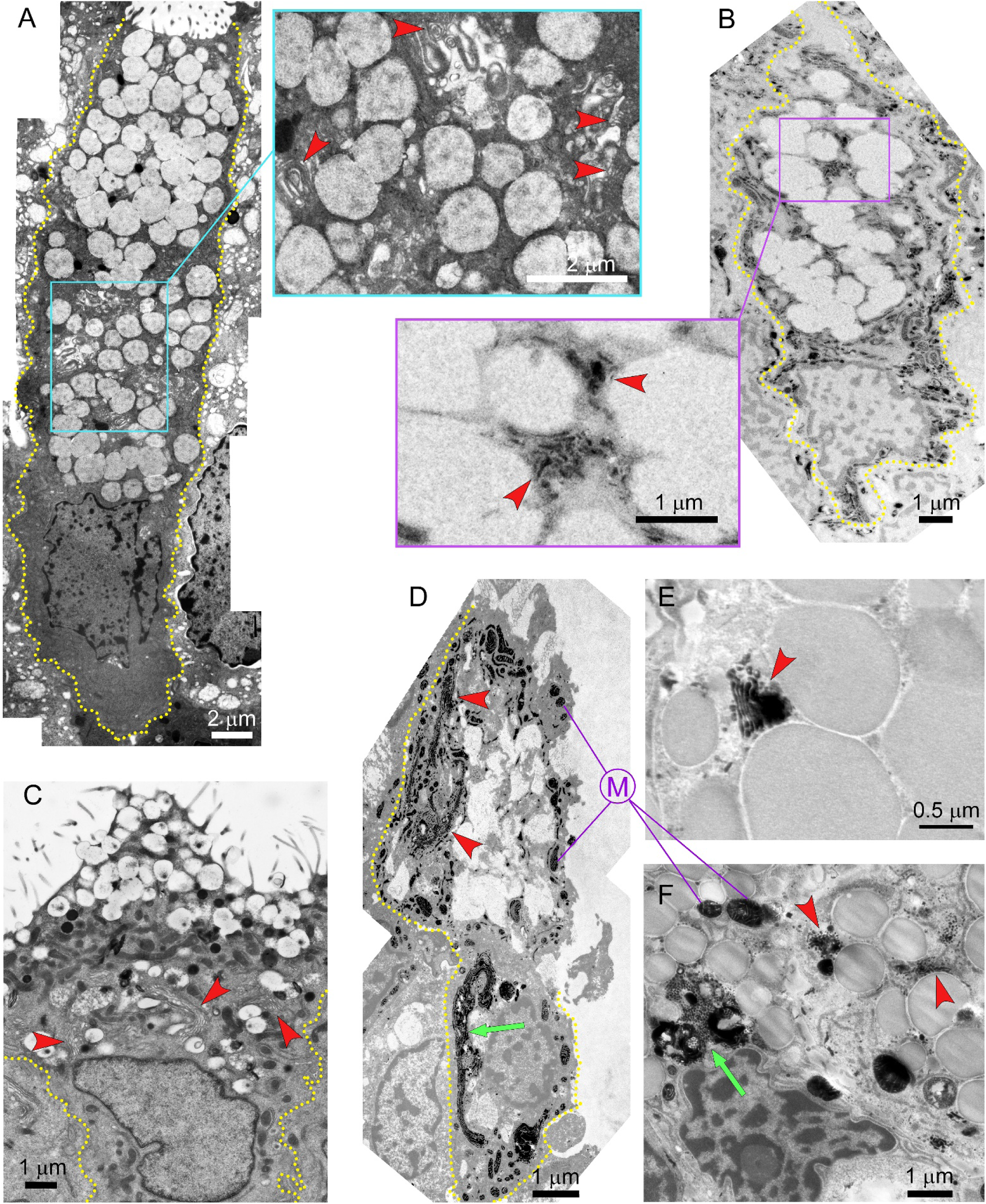
Electron microscopic images of Golgi structure in human airway secretory cells. *(A*) Profile of a human airway secretory cell compiled from multiple EM images of tissue fixed with glutaraldehyde, cacodylate and osmium tetroxide. The inset at high magnification shows multiple Golgi cisternae (red arrowheads) in close proximity to mucin granules*. (B)* Profile of a human airway secretory cell from EM images of tissue fixed and stained with zinc iodide – osmium tetroxide (ZIO) to highlight Golgi elements (red arrowheads) in close proximity to mucin granules. *(C)* EM profile of a cultured human airway epithelial secretory cell fixed with paraformaldehyde and glutaraldehyde, with Golgi cisternae (red arrowheads) seen both near the nucleus and in proximity to granules. *(D)* Partial EM profile of a cultured human airway epithelial secretory cell fixed and stained with ZIO, highlighting a perinuclear Golgi ribbon (green arrow) and peripheral Golgi satellites (red arrowheads), as well as stained mitochondria (purple M). *(E and F)* Golgi stacks (red arrowheads) adjacent to mucin granules in human submucosal gland mucous cells in tissue fixed and stained with ZIO, as well as a probable Golgi ribbon adjacent to a nucleus (green arrow in F).

### Golgi satellites are associated with immature mucin granules in human airway secretory cells

To determine whether Golgi satellites in human airway secretory cells are concentrated around immature mucin granules as in mice (Figures 1 and 2), we localized TGN46 with MUC5AC and MUC5B by immunofluorescence deconvolution microscopy. In proximal airways, tall secretory cells often contained large granules staining for MUC5AC, MUC5B, or both mucins together in the apical third of their cytoplasm (Figure 6A).

**Figure 6.**
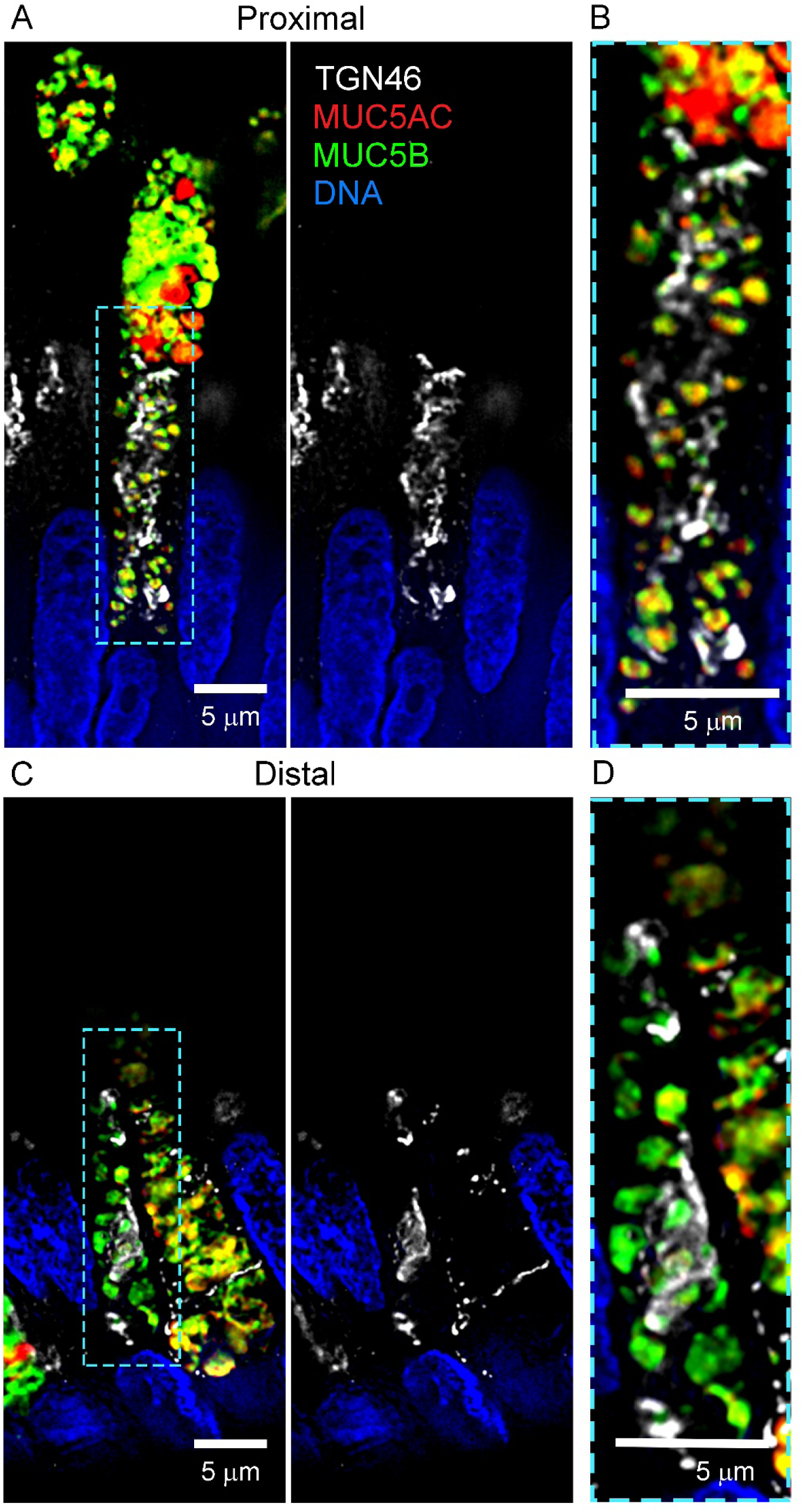
Localization of TGN in relation to mucin granules in human airway secretory cells. *(A)* Immunofluorescence deconvolution microscopy using antibodies against TGN46 to mark the trans-Golgi network, MUC5AC and MUC5B to mark mucin granules, and DAPI to mark nuclei in a proximal human airway. The MUC5AC and MUC5B channels are left out of the right image to better visualize TGN46. *(B)* The area marked in the middle of the secretory cell in *A* is shown at higher magnification. *(C)* Microscopy as in *A*, but in a distal human airway. *(D)* The area marked in the middle of the secretory cell in *C* is shown at higher magnification.

TGN46 was almost entirely excluded from this apical region containing mature mucin granules. In contrast, the middle third of these cells contained smaller, immature mucin granules with abundant TGN46 staining in puncta and tubules adjacent to the granules (Figure 6A-B). In distal airways, secretory cells contained more MUC5B than MUC5AC, and TGN46 staining extended to the apical plasma membrane (Figure 6C-D). Thus, proximal human airway secretory cells show TGN46 concentrated around immature mucin granules in the middle third of cells, similar to the distribution of TGN46 in metaplastic secretory cells in the axial bronchi of mice. Distal human airway secretory cells contained less MUC5AC than proximal cells, consistent with previous observations (23, 36). The distribution of TGN46 throughout the cytoplasm of distal secretory cells in close proximity to MUC5B granules suggests that these cells do not retain mature mucin granules, but instead continuously secrete them as previously suggested (20, 37). Additional immunofluorescence images of Golgi markers in relation to mucin granules are provided in Figure E8.

### Satellite TGN are a site of mucin packaging into granules

The concentration of Golgi satellites around immature mucin granules suggests that the presence of satellites in airway secretory cells is an adaptation for the synthesis and packaging of mucins. To address this, we examined the association of incompletely glycosylated mucin proteins (designated iMUC5AC and iMUC5B) in normal human tracheal tissue with mucin granules, ER, and Golgi satellites using monoclonal antibodies raised against a non-glycosylated peptide from MUC5AC (clone CLH2) (38) and almost completely deglycosylated purified MUC5B (clone PANH2) (39), neither of which reacts against fully *O*-glycosylated mature mucins. To label mature mucin granules, we used monoclonal antibodies that react with fully glycosylated and folded MUC5AC (40) or MUC5B (41) (designated fMUC5AC and fMUC5B). Mature mucin granules near the apex of secretory cells, identified either by the presence of fMUC5AC or fMUC5B (Figure E10A-B), did not show any reactivity with the antibodies against iMUC5AC and iMUC5B, which labeled small puncta in the middle third of cells. Some of the iMUC5AC and iMUC5B reactivity in smaller puncta overlapped with the ER marker calnexin (yellow color, Figure E10C-D), with nearby mucin granules appearing as black circles in those images. TGN46 immunofluorescence was observed adjacent to but not overlapping with puncta of iMUC5AC and iMUC5B (Figure E10E-F), suggesting that mucin glycosylation and folding are completed in adjacent satellite cisternae, and that the fully glycosylated mucins are then inserted into maturing granules at the TGN.

### Expression of Golgi satellites is upstream of mucin biosynthesis

Even though our findings suggest that the requirements of polymeric mucin synthesis and packaging drive the expression of Golgi satellites in airway secretory cells, the fact that the number of satellites changes minimally or not at all with mucous metaplasia (Figures E1, 1C and 2E) makes it unlikely that the synthesis of mucins itself drives satellite expression. To further address this, we examined secretory cells of mice in which both secreted polymeric mucins were deleted Fig. E11). The epithelium of double knockout mice appeared shorter than that of wild-type mice (Figure 7A-B), consistent with prior observations that airway epithelial height depends in part on the degree of mucin production (Figures 1, 2, E1, E2, E6, E7). However, there was no apparent reduction in the abundance of dispersed TGN46 puncta in secretory cells of double knockout mice compared to cells of wild-type mice (Figure 7A-B). Thus, we conclude that even though the unusual structure of the Golgi apparatus of airway secretory cells is likely an adaptation to facilitate the synthesis and packaging of mucins, expression of this structure is independent of actual mucin production and instead is determined during lineage specification.

**Figure 7.** Distribution of Golgi cisternae in mouse airway secretory cells lacking polymeric mucins. *(A)* Merged deconvolution immunofluorescence and differential interference contrast (DIC) images of the axial bronchus of a naïve wild-type mouse without mucous metaplasia. Antibodies against CCSP mark secretory cells, GM130 marks cis Golgi cisternae, TGN46 marks the trans-Golgi network, and DAPI marks nuclei. (*B*) The same field as in *A* but without CCSP staining or the DIC image to better show staining of Golgi cisternae. GM130-labeled cis Golgi cisternae localize near the nucleus in both secretory and ciliated cells, whereas the TGN46-labeled trans-Golgi network localizes alongside GM130 puncta in ciliated cells (yellow arrows) but is dispersed throughout the cytoplasm in secretory cells (yellow arrowheads). *(C)* Microscopy is as in *A*, but in MUC5AC and MUC5B double deletant mice. *(D)* The same field as in *C* but without CCSP staining or the DIC image. As in wild-type mice, TGN46 is dispersed throughout the cytoplasm only in secretory cells.

## DISCUSSION

Here we have analyzed the structure and function of the Golgi apparatus of airway secretory cells as a key intermediate in mucin production and secretion. We find that airway secretory cells express an unusual Golgi structure consisting of both a conventional perinuclear Golgi ribbon and numerous unconventional Golgi satellites. Previously, isolated Golgi stacks have occasionally been noted among mucin granules (42, 43), but no systematic characterization was performed. We find that satellites are de-enriched for the cis-Golgi marker GM130 relative to trans-Golgi and TGN markers. During mucous metaplasia, satellites redistribute from widespread dispersion throughout the cytoplasm towards concentration among immature mucin granules in the middle third of cells. Satellite number does not change with mucous metaplasia or with deletion of polymeric mucin genes. Several of these findings warrant further discussion.

The presence of dispersed Golgi stacks in various specialized vertebrate cell types is hypothesized to be driven by several different cell biological needs. These include local protein synthesis in neuronal dendrites to avoid the need for long distance transport if synthesis were to occur in the soma; serving as microtubule organizing centers for alignment of myofibrils in skeletal muscle cells; and the insertion of large secretory products into exocytic granules as occurs for von Willebrand factor (VWF) in endothelial cells (29–35). The most extensively studied of these systems is the neuron, where two types of dispersed Golgi stacks have been identified – outposts and satellites. Outposts occur proximally in dendrites where they function both in localized secretion as well as in microtubule nucleation related to dendritic branching (32, 44). Satellites occur distally in dendrites where they function in localized secretion of selected glycoproteins, are smaller than outposts, and lack cis-Golgi structural protein GM130. In view of the wide dispersion of Golgi stacks in airway secretory cells, their association with immature mucin granules (Figures 2, 5, 6, E7-8), their small size (average 0.19 x 0.68 µm, 4 cisternae; Figure 2), and their depletion of GM130 (Figures 3, 4, E6), we think it most appropriate to designate these dispersed Golgi stacks as satellites.

It is likely that the very large size of secretory mucin polymers has driven the adaptation of Golgi satellite expression in airway secretory cells, similar to the situation with VWF polymers that share homology with mucins. This inference is supported by the redistribution of satellites to close apposition to immature mucin granules in metaplastic cells engaging in a high rate of mucin synthesis (Figures 2, 5, 6, E8). By positioning satellites adjacent to nascent granules, mucin polymers could be directly transferred through tubules rather than requiring vesicular transport from a perinuclear ribbon, but this needs to be established experimentally. It is also possible that the advantage of concentrating specialized glycosyltransferases and chaperones required for mucin synthesis in a distinct Golgi compartment has driven the adaptation of satellite formation.

De-enrichment of secretory cell satellites for the cis-Golgi marker GM130 (Figures 3, 4, E6) raises the possibility that polymeric mucin synthesis begins in cis-Golgi cisternae in the ribbon, but is then transferred to satellites for further glycosylation and polymerization in medial and trans Golgi cisternae. Supporting this is possible continuity between the ribbon and some satellites in the perinuclear region of secretory cells (Figures E5, E8). However, the presence of abundant ER containing incompletely glycosylated mucins in the periphery of secretory cells (Figure E10) raises the alternative possibility that the ribbon is bypassed altogether for mucin synthesis, and that cis-Golgi enzymatic functions reside within satellites despite the absence of the structural protein GM130. In this scenario, the ribbon would instead be the site of synthesis of proteins other than mucins and of lipids. Consistent with biosynthetic roles for peripheral ER and satellites is the presence of numerous adjacent biosynthetic mitochondria (Figures 2, E3, E4). Atypical round mitochondria with few cristae have long been noted in airway secretory cells (43, 45). Recently, it was shown that mitochondria can partition between a classical elongated shape with prominent cristae that is specialized for energy production and a round shape lacking cristae that is specialized for biosynthesis (46, 47). Both types of mitochondria are observed in airway secretory cells, along with mitochondria with an intermediate phenotype (Figures 2A-B, E3, E4). Of interest, synthesis of proline, an abundant amino acid in mucins, is a primary function of biosynthetic mitochondria (46, 47).

There is a clear spatial separation between incompletely glycosylated mucins in the ER and fully glycosylated mucins in secretory granules (Figure E10A-D). The TGN lies immediately adjacent to puncta containing incompletely glycosylated mucins that likely are satellite Golgi cisternae (Figure E10E-F). This suggests that the TGN itself is not a site of final mucin synthesis, but rather a site of packaging of fully glycosylated mucins into granules, consistent both with classical views of TGN packaging function (27, 28) and the recent identification of a sorting function for TGN46 itself (48, 49). Possible signals on mucins of the completion of biosynthesis and readiness for packaging might be the addition of terminal sugars such as sialic acid or fucose, or modifications such as sulfation. Whether enlargement of mucin granules in the progression from immature to mature granules (Figures 2A-B, E4) occurs exclusively due to ongoing transfer of mucins from satellite TGN to maturing granules or is also due to lateral fusion between immature granules is not known.

While it is likely that the need to synthesize and package polymeric mucins has driven the expression of satellites in airway secretory cells, it is striking that the number of satellites does not rise substantially with increased mucin synthesis during mucous metaplasia (Figures 1, 2, E1). The small increases observed by immunofluorescence microscopy could be due to cellular expansion during metaplasia allowing better visualization of fluorescent puncta (Figures 1, E1), and the lack of increase observed by EM supports that interpretation (Figure 2). Even more striking, the number of satellites does not fall with absent mucin synthesis in double deletant mice (Figure 7). Thus, expression of satellites is apparently specified during secretory cell development and remains relatively constant independent of polymeric mucin synthesis. This differs from the expression of satellites in neurons that depends on neuronal activity (44).

Furthermore, there is no substantial change in the size of satellites during secretory mucous metaplasia (Figure 2). The capacity of pre-existing satellites to meet increased mucin synthetic demand is reminiscent of the lack of increase in the abundance of the exocytic protein Munc18b during mucous metaplasia (50). In view of the very small number of mucin granules in non-metaplastic secretory cells (Figures 2, E3), it would be interesting to determine whether the abundant Golgi satellites also perform synthetic or packaging functions for proteins other than mucins.

The Golgi apparatus is increasingly recognized as being a site of sensing and scaffolding for numerous cellular functions including cytoskeletal organization, sensing of proteostasis stress, metabolism, autophagy, inflammation, and apoptosis (29, 31). Furthermore, some of these cellular functions have been shown to vary with changes in Golgi structure. In view of the postulated roles of many of these cellular functions in lung pathobiology, gaining further knowledge of Golgi apparatus structure and function in airway epithelial secretory cells that are constantly sensing and responding to the external environment and to immune signals seems warranted.

**Author disclosures** are available with the text of this article at www.atsjournals.org.

## Acknowledgement

We are indebted to Peter A. Nielsen and Henrik Clausen for the PANH2 and CLH2 antibodies, and to Sergio Trillo-Muyo for the anti-MUC5B antiserum. The content is solely the responsibility of the authors and does not represent the official views of the Department of Veterans Affairs or the United States government.

## Supplemental Materials and Methods

### Materials

All chemicals and supplies were purchased from MilliporeSigma unless indicated otherwise.

### Mice

C57BL/6J mice of both sexes were purchased from the Jackson Laboratory and bred to form cohorts. We have found no difference in mucous metaplasia or secretion efficiency between female and male mice from ages 6 to 26 weeks (1), so mice of both sexes within this age interval were used. Mice were housed up to 5 per cage in individually ventilated cages in a specific pathogen-free facility where the exposure to dust was minimized by quarter-inch corn cob bedding, in controlled temperature (21°C) and relative humidity (55%), and a 12-hour light/dark cycle. Standard chow and water were available ad libitum. The animal care and experimental protocols were approved by the Institutional Animal Care and Use Committee of MD Anderson Cancer Center (MDACC) (permit number 00001214-RN02).

For induction of mucous metaplasia, a single dose of 1-3 µg mouse recombinant IL-13 (BioLegend) in 40 µl PBS was instilled into the posterior pharynx by direct visualization under isoflurane anesthesia, which mice then aspirated into their lungs. Lungs were harvested 72 h after the cytokine doses under tribromoethanol anesthesia, inflated at 20 cm water pressure with 4% neutral buffered paraformaldehyde for 24 h at 4°C overnight, dehydrated with graded ethanol, cleared with Histo-Clear (HS-200, National Diagnostics), then embedded in paraffin. A single transverse 5-µm section of the axial bronchus of the left lung was taken from each mouse between the first and second lateral branches as described (1).

For Figure 7, we used tissue from *Muc5ac/Muc5b* double deletant mice that were generated using CRISPR/NHEJ at the University of Colorado (IACUC permit number 46). Gene disruption was achieved by deleting exon 1 as was done previously in embryonic stem cells for both *Muc5ac* (2) and *Muc5b* (3) individually (Fig. E11A). Here, *Muc5ac*/*Muc5b* double knockout mice were produced by targeting intact *Muc5ac* alleles in *Muc5b*^lox/lox^ mice. Guide RNAs 5’-GCTAGTCAATGGCAGTAGTC and 5’-GGCTGGGCATCCAATGTGTG were mixed with Cas9, sperm from *Muc5b*^lox/lox^ males, and oocytes from C57BL/6 females. The mixture was used to perform in vitro fertilization by Dr. Jennifer Matsuda at National Jewish Health.

F0 pups were screened for *Muc5ac* disruption by PCR using primers 5’-AGCTCAGGGAGAGTCTCAAA (forward) and 5’-AGGCTAAGAGAAACCACATTCC (reverse). Wild type mice yielded an 841 bp amplicon, and mice that had undergone CRISPR with non-homologous end-joining yielded a 332 bp fragment. Sequencing confirmed this to have successfully excised exon 1 from the *Muc5ac* gene. Subsequent breeding confirmed disruption of *Muc5ac* in cis with *Muc5b*^lox^ alleles. *Muc5ac*^+/-^;*Muc5b*^lox/+^ mice were then crossed with CMV-Cre transgenic mice (B6.C-Tg(CMV-cre)1Cgn/J, Strain #006054, Jackson Laboratory) to produce *Muc5b* knockout alleles in all tissues, including the germline. Heterozygotes also carrying Cre transgenes were crossed with C57BL/6J mice (Jackson Laboratory), and resulting *Muc5ac*^+/-^; *Muc5b*^+/-^ mice were bred again with C57BL/6J mice to confirm germline inheritance independent of continued CMV-Cre transgene presence. Confirmation of *Muc5ac* gene disruption was validated using RT-qPCR on stomach tissues, which demonstrated high levels of baseline *Muc5ac* transcripts in *Muc5ac*^+/+^;*Muc5b*^lox/lox^ mice and *Muc5ac* mRNA absence in *Muc5ac*^-/-^; *Muc5b*^-/-^ littermates (Fig. E11B).

*Muc5ac*^+/-^;*Muc5b*^+/-^ mice were bred together to produce experimental animals used here. Notably, there was a strong effect of double deficiency that was apparent at all stages of life. Due to their linkage on mouse chromosome 7, intercrossing *Muc5ac*^+/-^;*Muc5b*^+/-^ mice was expected to yield simple Mendelian inheritance ratios. Double knockout pups were observed born at only 8.1% frequencies (*p* < 0.0001, chi square). By contrast, *Muc5ac* and *Muc5b* single knockout lines produced homozygous deletants at 23% and 21% frequencies (*p* = 0.136 and 0.054, respectively). In addition to these observations in early life, there were also significant effects of combined *Muc5ac* and *Muc5b* gene deficiency on post-natal survival. Tracking cohorts of mice over one year revealed significant early mortality in single *Muc5b* knockout mice, consistent with what was reported previously (3), and deficiency in both mucins caused even greater survival impairment (Fig. E11C).

Median survival in *Muc5ac*^-/-^; *Muc5b*^-/-^ mice was only 6.6 weeks, which was significantly lower than 32.6 weeks median survival in *Muc5b*^-/-^ mice (*p* = 0.0004).

### Human tissue

For examination of most human airways, de-identified tissue from lungs donated but not used for transplantation was obtained by the Pulmonary Center of Excellence Biobank at the University of Texas Health Science Center at Houston under an approved institutional review board protocol (HSC-MS-08-0354). None of the donors had known lung disease (Table S1). For proximal airways, bronchial tissue was dissected free from major vessels and pleural tissue. A 1-1.5 cm longitudinal piece of lobar or segmental bronchus near the midpoint between its take-off and its termination was excised and placed in 10% neutral buffered formalin for 24 h at 4°C overnight, then embedded in paraffin. For distal airways, a section was made in the peripheral lung parallel to the pleural surface, 1-2 cm from the pleura, then fixed and embedded as above. Airways 200-1,000 µm in diameter and lacking submucosal glands were then selected for microscopic analysis at MDACC.

For the imaging of immature mucins in Figure E10, normal human tracheal tissue was obtained from transplant donor tissue collected at the Sahlgrenska University Hospital, Gothenburg, Sweden following protocols approved by the Swedish Ethical Review Authority (Etikprövningsmyndigheten, 2020-02658). The tissue was fixed overnight in 4% neutral buffered formalin and then embedded in paraffin.

### Immunofluorescence microscopy of mouse and human airways

For imaging of mouse and human airways at MDACC, paraformaldehyde-fixed paraffin-embedded tissue blocks were cut into sections 5 μm thick, deparaffinized with xylene, and rehydrated with ethanol/water. Samples were then placed in a 10 mM sodium citrate bath (pH 6.0) and heated in a pressure cooker for 10 min. After cooling, lung sections were washed with PBS and permeabilized with 0.3% Triton X-100 in PBS for 20 min. Sections were then blocked with 5% normal donkey serum (NDS) in PBS and 0.05% Tween 20 (PBST) for 1 h at room temperature. Following PBS and PBST washes, background autofluorescence was quenched with an electrostatic agent (Vector TrueVIEW) for 3 min. Sections were given a final PBS wash and mounted with Fluorescence Mounting Medium (Dako). Antibodies used for immunofluorescence microscopy are listed in Table S2.

For enumeration of TGN elements, a pilot study was performed using a confocal microscope (Nikon A1, Japan) with a 60x lens (Apochromat TIRF 60XO, NA 1.49, Nikon) with oil immersion (Figure E1). Settings were optimized using the Nyquist rate. Z-stacked optical sections were evaluated for TGN46 puncta in 0.5 μm steps to avoid counting the same structure twice, and cell borders were determined from differential interference contrast images. For all subsequent studies except Figure E10, widefield fluorescence microscopy with post-acquisition deconvolution of the images was performed as follows. Z-stacked images of 0.5 μm steps were taken on a DeltaVision Elite microscope (GE Healthcare) using a 100x oil immersion objective (UPLSAPO 100XO 1-U2B836, NA 1.4, Olympus) and 0.23 µm optical sections. Images were then deconvolved with SoftWoRx software (GE Healthcare). For enumeration of TGN elements (Figure 1C), TGN46 puncta were counted on optical sections 1 µm apart.

For imaging of immature mucins in human tissue at the University of Gothenburg, formalin-fixed, paraffin-embedded tissue blocks were cut into sections 6 µm thick, deparaffinized with xylene, rehydrated in a series of graded ethanol baths, followed by antigen retrieval in 10 mM citrate buffer (pH 6.0). Following this, sections were permeabilized with 0.1% Triton X-100 in TBS for 10 min and then blocked with 1% bovine serum albumin/10% donkey serum in TBS for 1 h at room temperature. Afterwards, sections were incubated sequentially with primary antibodies raised against mature MUC5AC (clone 45M1), mature hMUC5B (MUC5B-D3 antibodies), incompletely glycosylated MUC5AC (iMUC5AC, clone CLH2), incompletely glycosylated MUC5B (iMUC5B, clone PANH2), TGN46, and calnexin as detailed in Table S2. Antibodies against iMUC5AC were raised against a non-glycosylated hMUC5AC peptide, and antibodies against iMUC5B were raised against purified hMUC5B deglycosylated with trifluoromethanesulfonic acid. Following incubation with primary antibodies, sections were washed and incubated with appropriate fluorophore-conjugated secondary antibodies as detailed in Table S2. Cell nuclei were counterstained with Hoechst 34580 (5 µg/ml in TBS). High resolution confocal micrographs were acquired on an upright LSM900 confocal microscope equipped with an Airyscan2 detector (Carl Zeiss) using a Pan-Apochromat 63x/1.4 oil DIC M27 lens. Confocal micrographs were captured using Zen software (Zen blue v3.1; Carl Zeiss), data was exported to Imaris (v9.5; Bitplane), and images were exported in TIFF format.

### Transmission electron microscopy

For studies in mice, mild mucous metaplasia was induced with a single dose of 0.2 μg IL-13 instilled intrapharyngeally as described above for immunofluorescence studies, then mice were anesthetized and sacrificed five days later. Lungs were excised and fixed in 2.5% glutaraldehyde in 0.1 M sodium cacodylate buffer (pH 7.2) containing 20 mM calcium chloride for 2 h, followed by a 1-h secondary fixation in buffered 1% osmium tetroxide. The fixed left lung was then sectioned into a single transverse cut of the axial airway between lateral branches 1 and 2 and embedded in EMbed 812 epoxy resin (14120, Electron Microscopy Sciences). Sections 100 nm thick were stained with uranyl acetate and lead citrate and were viewed in a Tecnai 12 transmission electron microscope. Secretory airway epithelial cells were identified by an apical plasma membrane contacting the airway lumen, the presence of secretory granules, and absence of cilia. Cells were selected for further imaging if a complete plasma membrane was visible that contacted the airway lumen, neighboring cells on both sides, and the interstitium on the basal surface, and if a portion of the nucleus at least 2 µm wide was visible. Multiple photographs of each secretory cell were assembled using Adobe Photoshop into photomontages containing the complete cell. These composite images, termed “cellular profiles”, were then analyzed for the distribution of Golgi elements, which were identified as stacks of membrane-bound cisternae (8). To measure the position of a satellite on the apical-basal axis, a line from the middle of the nucleus to the middle of the apical membrane (NA) was set as +1, and the distance from the middle of the nucleus to the satellite (NS) was calculated as a fraction (Figure 2C).

Satellites below the middle of the nucleus were calculated as negative fractions. To measure the position of a satellite on the lateral axis, a line from the central apical-basal axis of the cell to the lateral membrane (CL) was set as +1, and the distance from the central apical-basal axis to the satellite (CS) was calculated as a fraction (Figure 2D).

For routine studies of human tissue, fixation and staining were as described above for mouse tissue. To highlight Golgi elements in some studies, glutaraldehyde-fixed tissue was stained with Zinc Iodide-Osmium tetroxide reagent (ZIO) according to (9), with minor deviations. Briefly, 3 g of Zn powder (Sigma #243469) were suspended in 20 ml of water, then 3 g of I_2_ flakes (Sigma # 376558) were added gradually with vigorous mixing to prevent overheating. The suspension was then filtered, and 4 ml of the colorless solution were mixed with 2 ml 50 mM Tris (pH 7.4) and 2 ml 2% OsO_4_ in water, resulting in a light brown solution. Tissue samples were immersed in this solution for more than 100 h in the dark at 4°C. For studies in cultured human airway epithelial cells at both the University of Nebraska (Figure 5C) and MDACC (Figure 5D-F), non-diseased human airway epithelial cells (HAECs) were derived for culture from excess airway tissue donated for lung transplantation as previously described (10). After 5–7 days of expansion with BEGM media (Lonza #CC-3170), air–liquid interface (ALI) conditions were established and HAECs were fed thereafter with PneumaCult ALI media. Beginning 14 days after the establishment of ALI, cells were treated 7 days with IL-13 (10 ng/mL), then fixed. Cells cultured at the University of Nebraska were fixed with 2% paraformaldehyde and 2.5% glutaraldehyde in 100 mM cacodylate buffer (pH 7.2) for 2 h at room temperature, followed by a 1-h secondary fixation in buffered 1% osmium tetroxide. They were viewed on a JEOL 1200 EX electron microscope equipped with an AMT 8 megapixel digital camera (Advanced Microscopy Techniques). Cells cultured at MDACC (Figure 5D-F) were stained with ZIO and viewed on a Tecnai 12 electron microscope as above.

### Serial block face – scanning electron microscopy (SBF-SEM)

This was performed on mouse tissue with mild mucous metaplasia induced with a single dose of 0.2 μg IL-13 instilled intrapharyngeally as described above. Tissue was fixed as for transmission EM. SBF-SEM was performed using a 3View®2 system (Gatan) mounted in a MIRA3 field emission scanning electron microscopy (TESCAN) as previously described (11). In brief, serial imaging was conducted under high vacuum using a Schottky emitter and an accelerating voltage of 9 kV with spot size of 4.3 nm and pixel size ranging from 2.7-5.9 nm. Backscatter electron detection was used to image the block face. Serial images were acquired at 100 nm intervals. Image stacks were post-processed for spatial drift removal using Gatan DigitalMicrograph software. Segmentation and reconstruction of the Golgi apparatus from a secretory cell was performed using Amira 6.0.1 software (FEI Systems) as previously described (12). In brief, the nucleus was reconstructed along its surrounding nuclear envelope, the Golgi apparatus by both its electron dense staining and cisternal processes, and the associated secretory granule along its limiting membrane based on the electron density of its lumenal content as described in the legend to Figure E5G.

## Data Analysis

Statistical analyses were performed and plots generated using GraphPad Prism version 8.4.3 (Prism software), with *P* < 0.05 considered significant. Methods of analysis, *P* values and *n* values for each sample are included in figure legends.

## Supplemental Results

**Calculation of the number of Golgi satellites per mouse airway secretory cell visible by electron microscopy** (related to Figure 2E). From a sample of EM cellular profiles of both naïve and metaplastic airway secretory cells (five each), we found an average cell width of 10.4 µm and height of 16 µm. Assuming that a cell has a cylindrical shape, the average cross-sectional area (3.14 x radius 5.2 µm^2^) is 84.9 µm^2^, and volume (84.9 µm^2^ x height 16 µm) is 1,358 µm^3^. Further assuming that an EM image is roughly vertical on an apical-basal axis, passes through the center of a cross-sectional circle, and is seen en face, the average cross-sectional area (width 10.4 µm x height 16 µm) is 166.4 µm^2^, and volume (area 166.4 µm^2^ x section thickness 0.1 µm) is 16.6 µm^3^. Based on these assumptions, an EM image would sample 16.6 µm^3^/1,358 µm^3^ (0.012%) of the cell volume. With a mean of 0.68 satellites visualized in naïve cells per EM image, the mean total satellite number per cell is estimated at 55.5, and with a mean of 0.71 satellites visualized in metaplastic cells, the mean total satellite number per cell is estimated at 57.9.

## Supplemental Figure Legends

**Figure E1.**
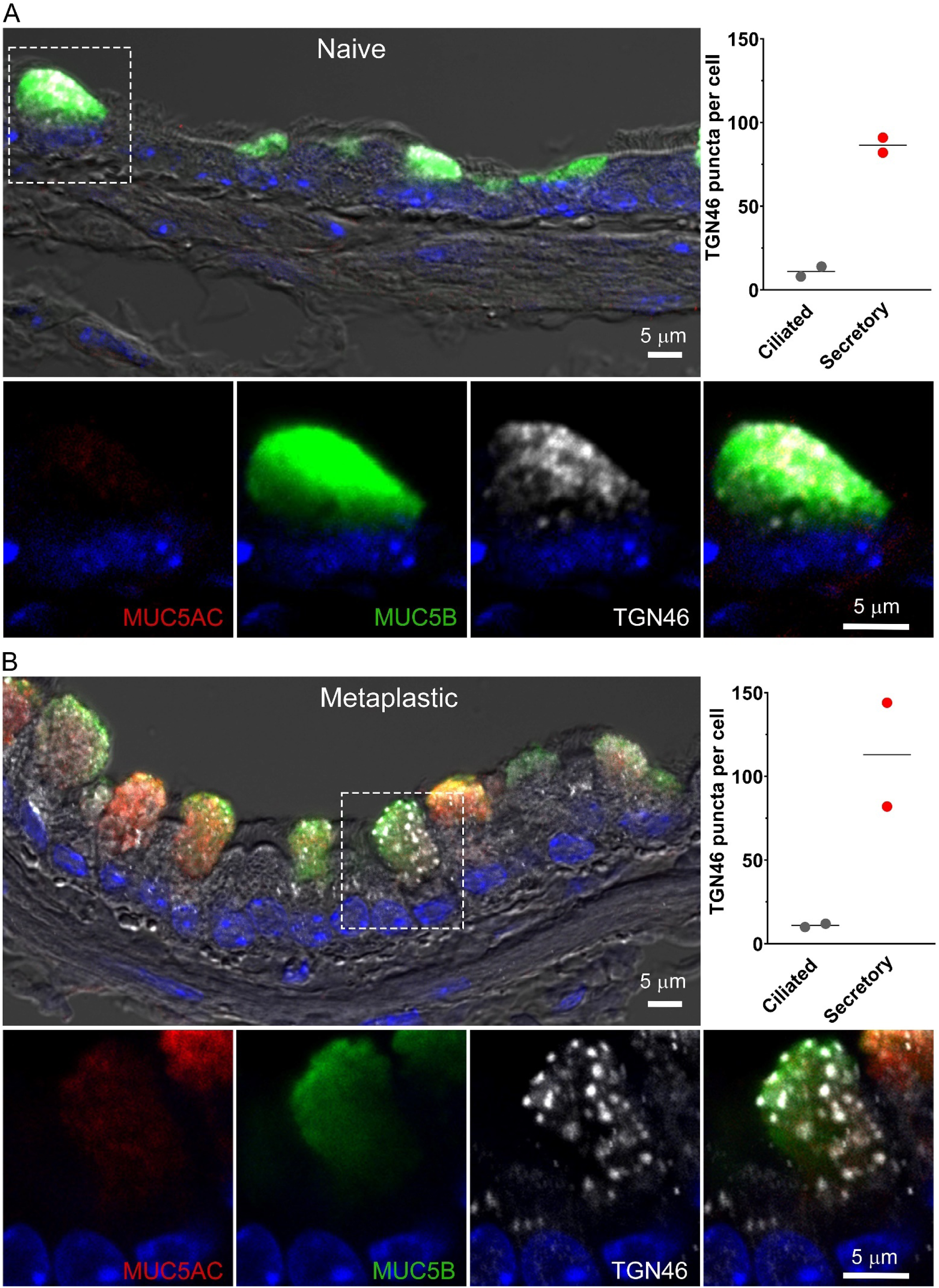
Numerous dispersed TGN elements in mouse airway secretory cells by confocal immunofluorescence microscopy. Laser confocal micrographs of mouse airways stained with antibodies against MUC5AC, MUC5B, and TGN46, with nuclei stained with DAPI, and merged with differential interference contrast (DIC) images. (*A*) Naïve, uninflamed airway with no visible MUC5AC staining. Secretory cells are marked by MUC5B staining. Dashed box outline in top left image shows region of interest magnified in dual channel fluorescence views without DIC in left three panels of bottom row. The total number of TGN46 in two ciliated and two secretory cells, measured as described in Methods, is shown in the top right panel. (*B*) Airway with mucous metaplasia due to IL-13 instillation showing MUC5AC staining. Layout of the panels is as in *A*.

**Figure E2.**
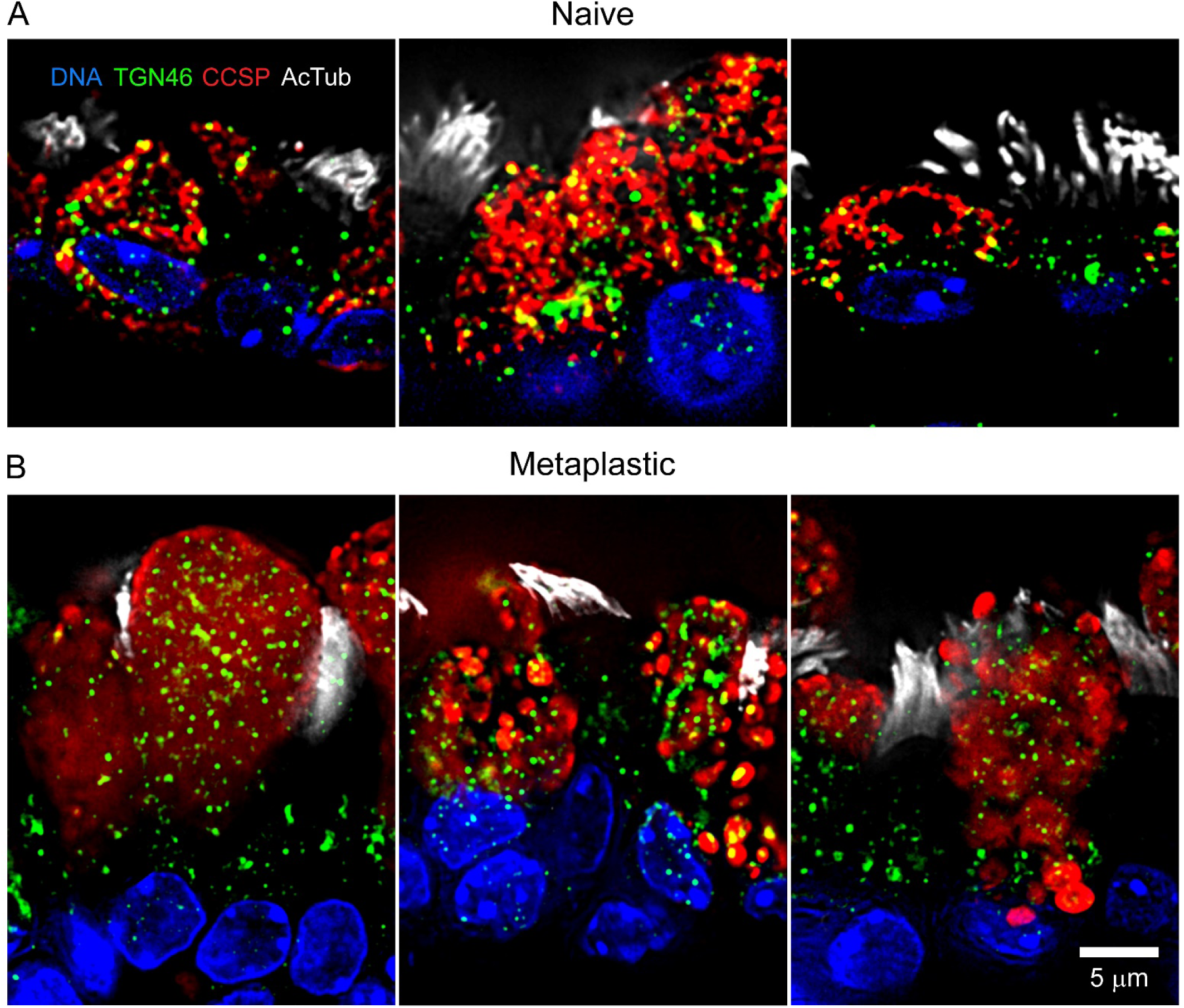
Additional images of dispersed TGN in mouse airway secretory cells by deconvolution immunofluorescence microscopy. Immunostaining is exactly as in Figure 1 using antibodies against CCSP to mark secretory cells, acetylated tubulin (AcTub) to mark ciliated cells, TGN46 to mark the trans-Golgi network, and DAPI to stain nuclei in the axial bronchi of mice. (*A*) Three naïve airways without mucous metaplasia. (*B*) Three airways with mucous metaplasia due to IL-13 instillation.

**Figure E3.**
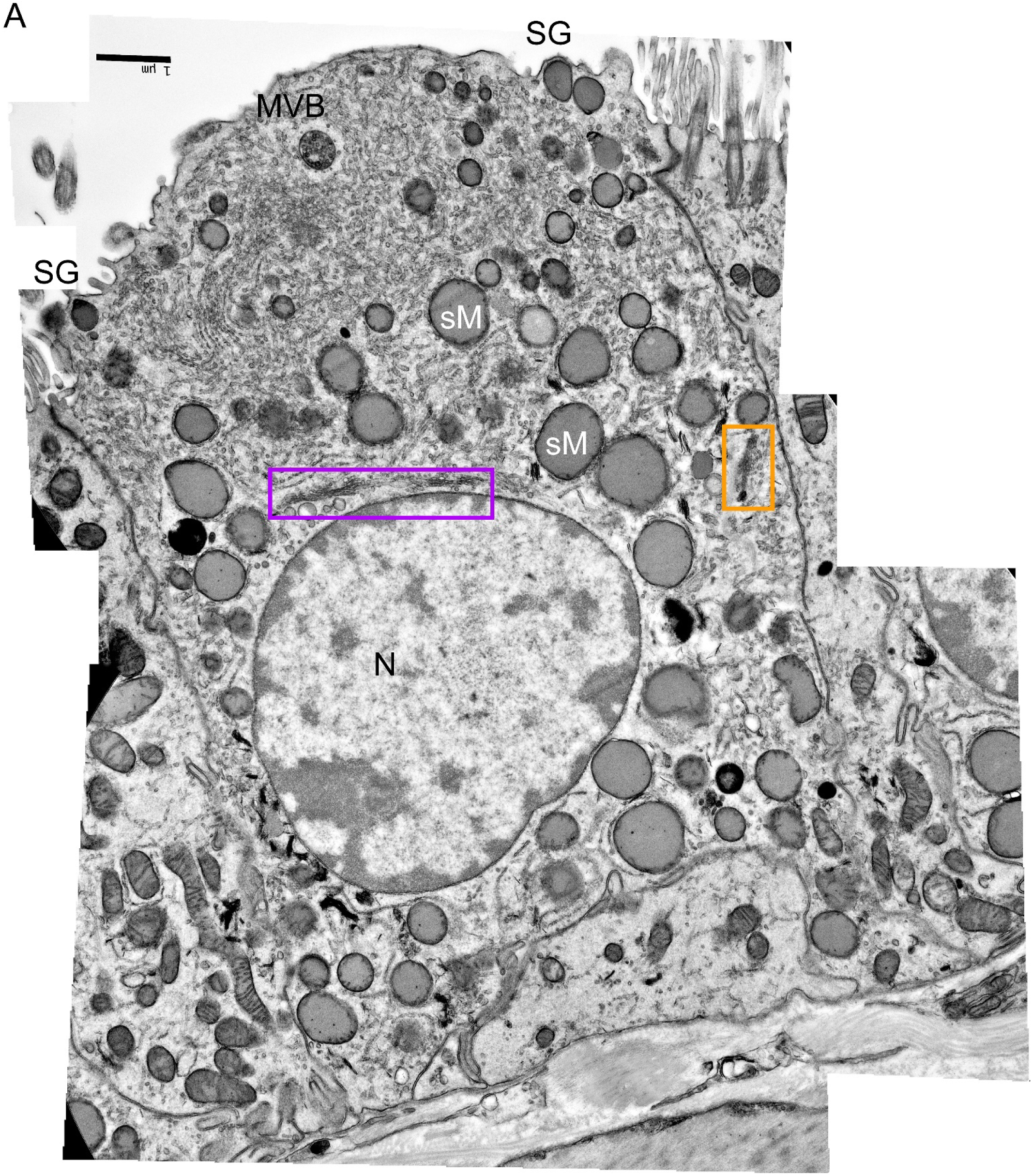
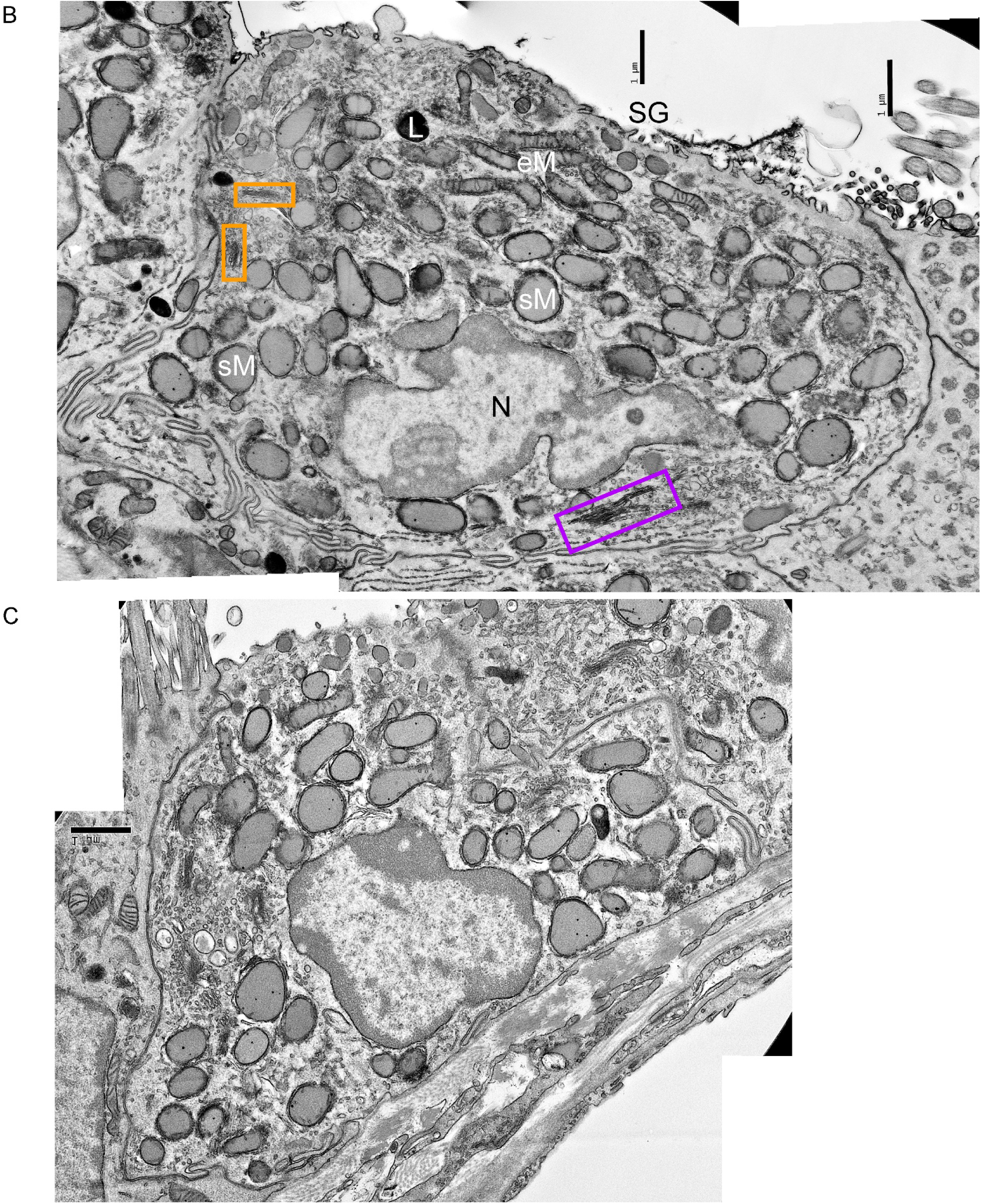
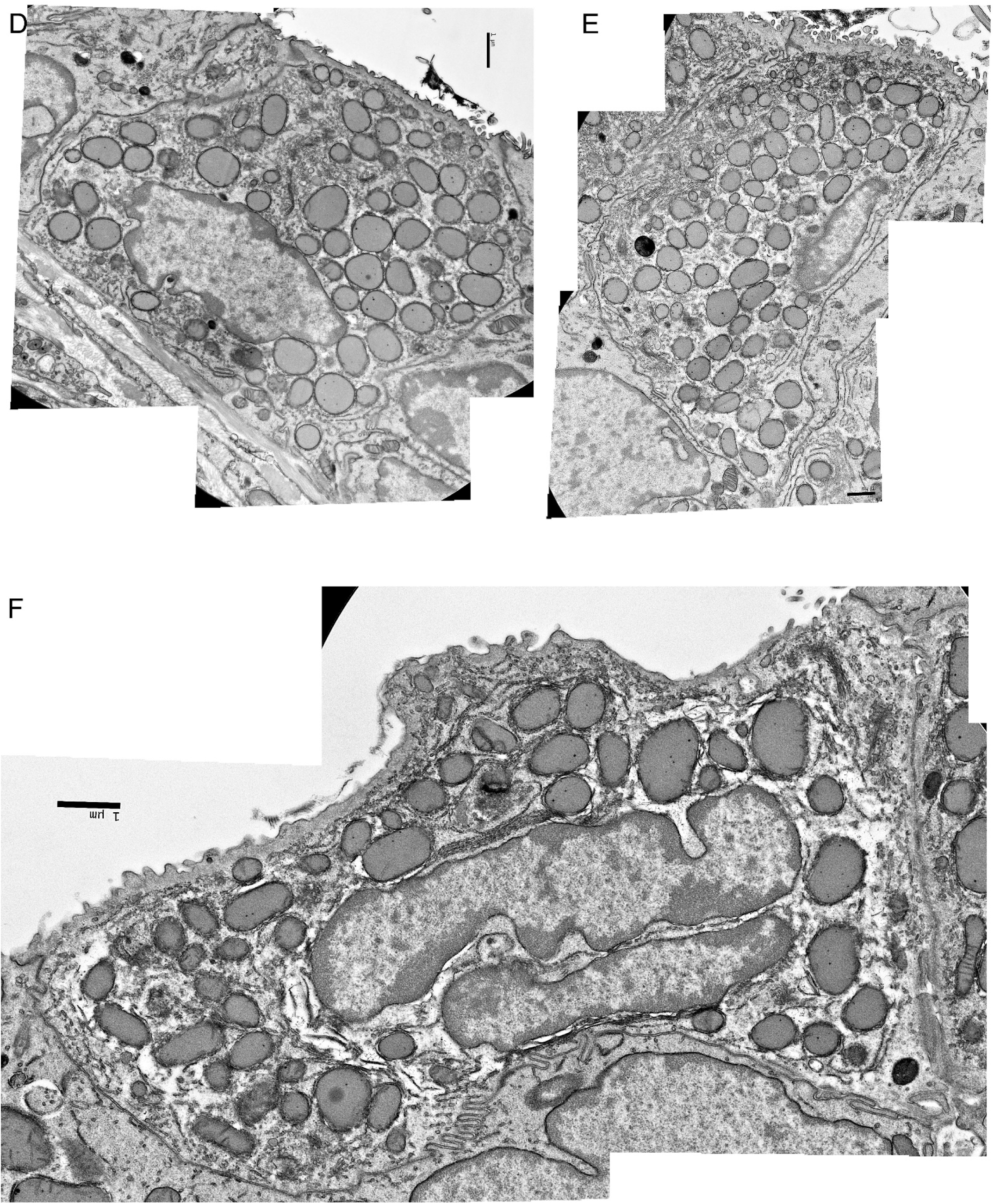
Additional EM images of Golgi satellites and ribbons in naïve mouse airway secretory cells. Cell profiles compiled from multiple electron micrographs of secretory cells as in Figure 2. (*A*) The same cellular profile shown in Figure 2A is enlarged and annotated here. The Golgi ribbon is outlined in magenta and a Golgi satellite is outlined in orange. SG = secretory granule, MVB = multivesicular body, eM = mitochondrion specialized for energy production, sM = mitochondrion specialized for synthesis, iM = mitochondrion intermediate between energy and synthesis specialization, L = lysosome, N = nucleus. (*B*) Another secretory cell profile annotated as in *A*. (*C-E*) Additional secretory cell profiles not annotated.

**Figure E4.**
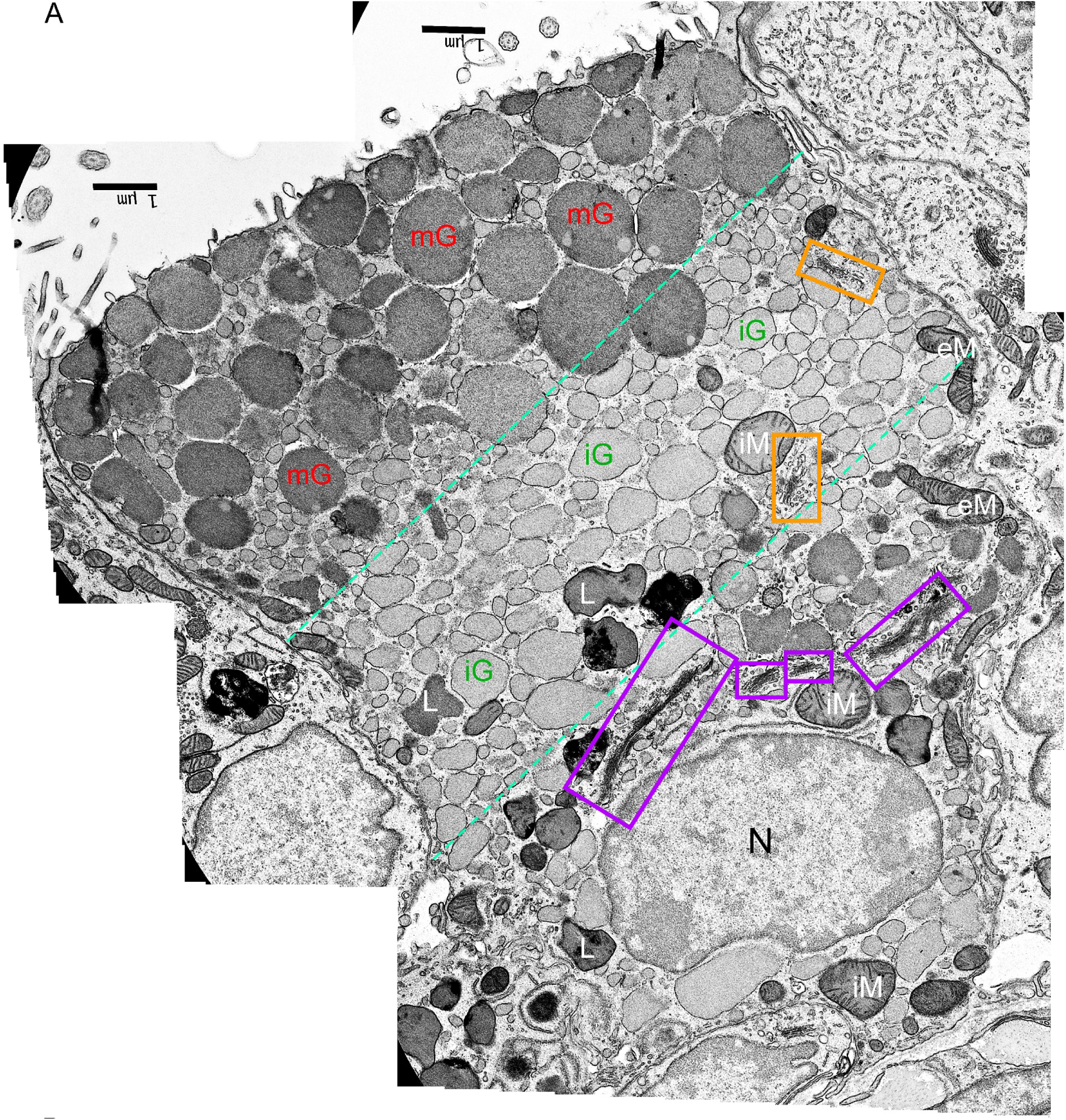
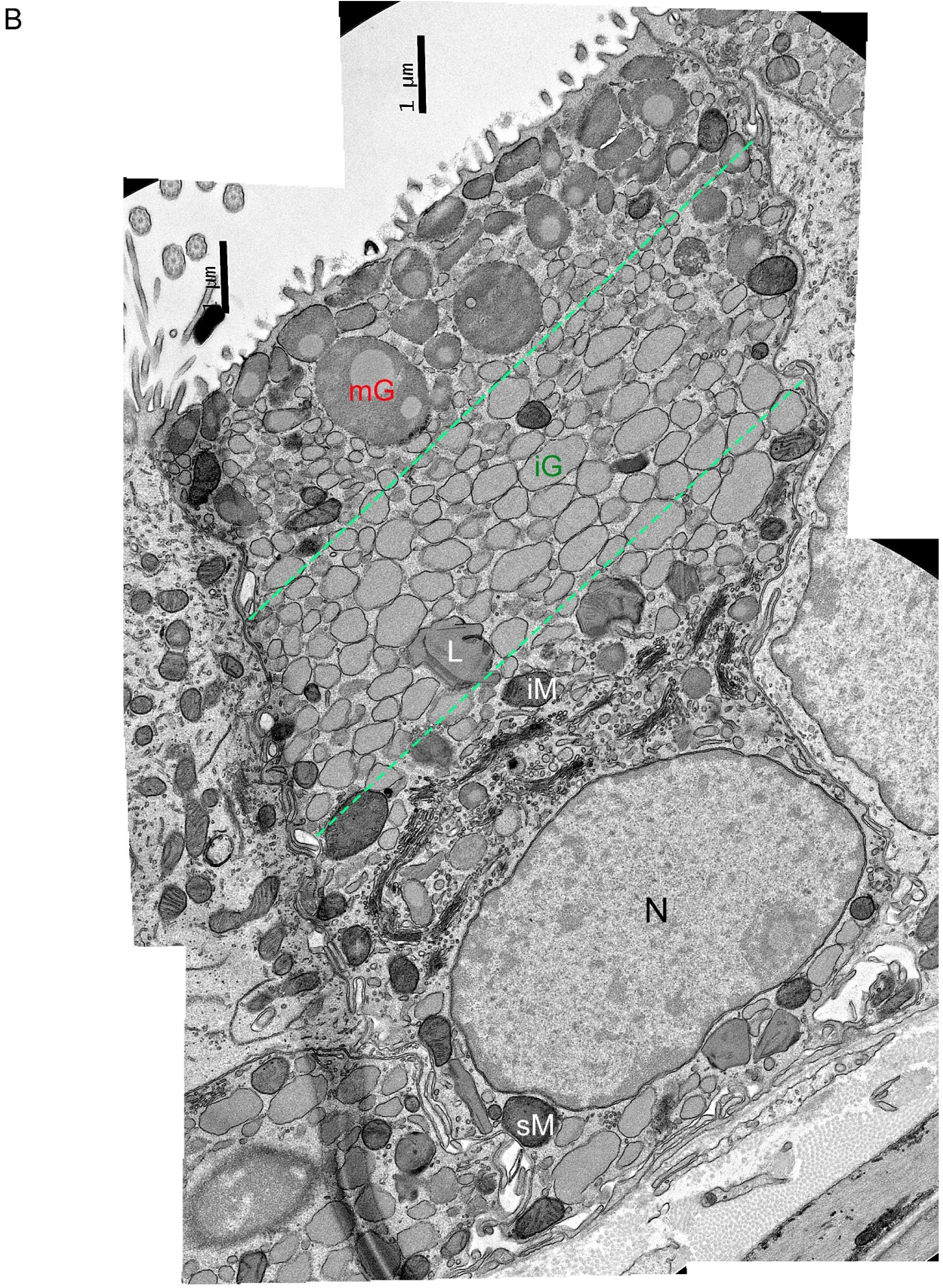
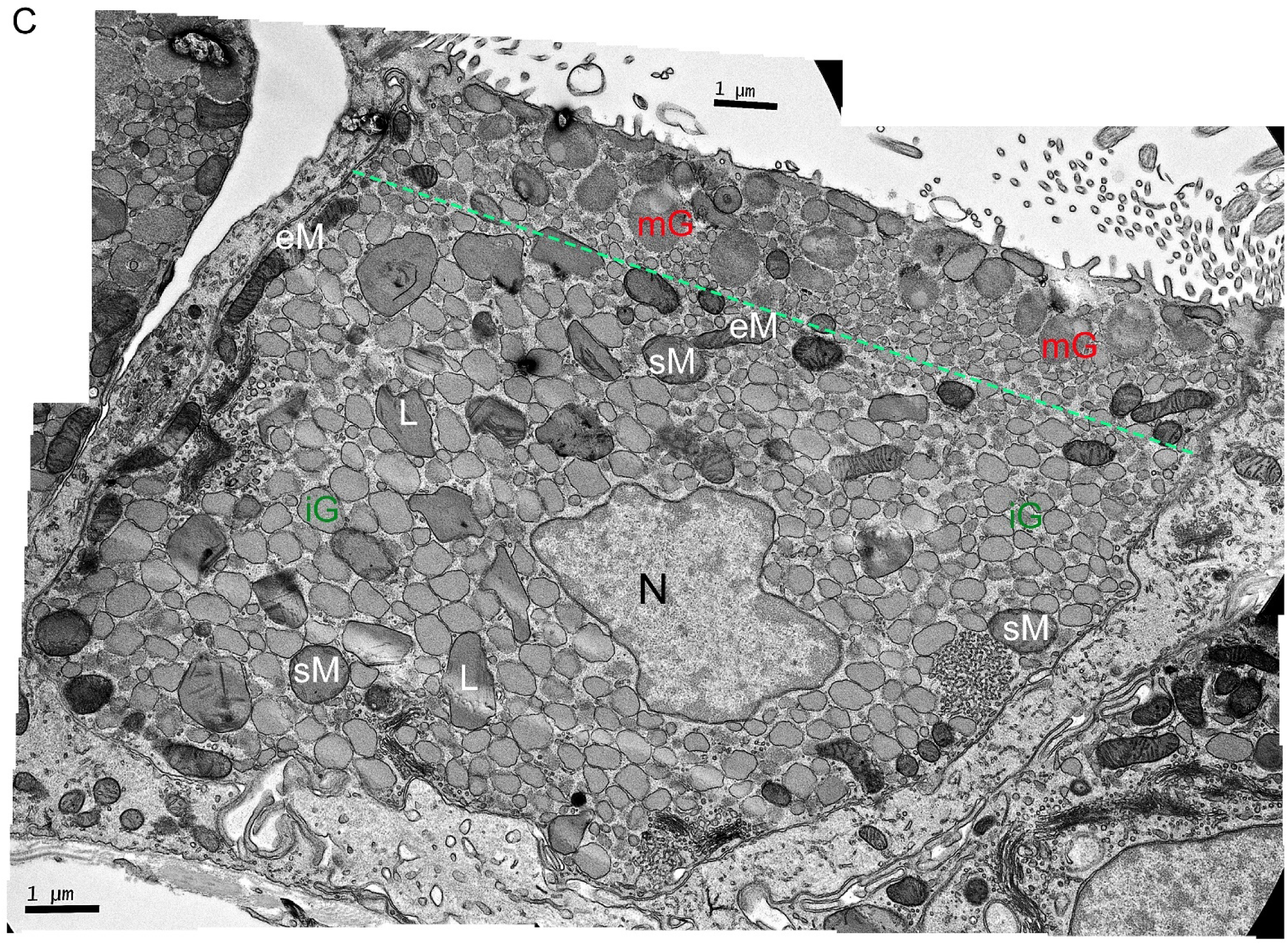
Additional EM images of Golgi satellites and ribbons in metaplastic mouse airway secretory cells. Cell profiles compiled from multiple electron micrographs of secretory cells as in Figure 2. (*A*) The same cellular profile shown in Figure 2B is enlarged and annotated here. The Golgi ribbon is outlined in magenta and two Golgi satellites are outlined in orange. The dashed teal lines mark boundaries between the apical third of the cell filled with mature mucin granules, the middle third filled with immature mucin granules, and the basal third mostly devoid of mucin granules. Mature mucin granules are larger than immature granules, more electron dense, rounder, and often contain a small spherical region of differing electron density that is uniformly observed in serial block face – scanning electron microscopy (SBF-SEM) (red arrowhead in Figure E5J-L; Video 2). mG = mature mucin granule, iG = immature mucin granule, eM = mitochondrion specialized for energy production, sM = mitochondrion specialized for synthesis, iM = mitochondrion intermediate between energy and synthesis specialization, L = lysosome, N = nucleus. (*B*) Another secretory cell profile partially annotated as in *A*. (*C*) An additional secretory cell profile annotated as in *A*, but with only a single boundary marked between the apical pole containing mature mucin granules and the middle region of the cell filled with immature mucin granules and mitochondria.

**Figure E5.**
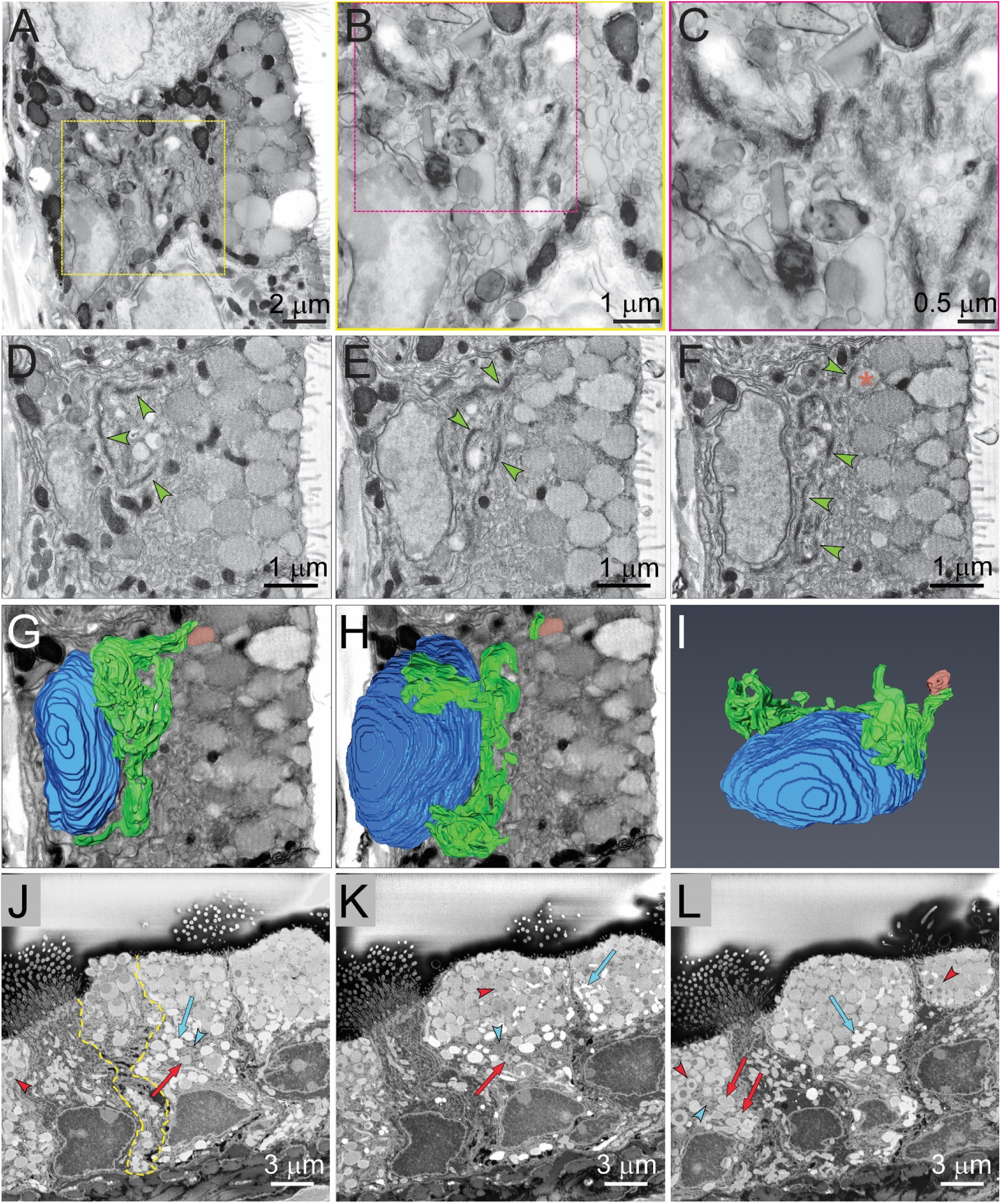
Serial block face – scanning electron microscopy (SBF-SEM) of metaplastic mouse airway epithelial cells. Tissue from mice with mild mucous metaplasia induced by instillation of IL-13 into the airway was fixed and processed for SBF-SEM as described in Methods. Videos were constructed from the serial scanning EM images, with panels *A-F* taken from Video 1 and panels *J-L* from Video 2. (*A*) A high resolution image showing Golgi stacks, mucin granules, mitochondria, and portions of two nuclei. (*B*) The boxed area in A is shown at higher magnification. (*C*) The boxed area in *B* is shown at higher magnification. (*D*) An image corresponding to Video 1 at 7 seconds showing a Golgi ribbon (green arrowheads) just above the cell’s nucleus (left). (*E*) An image from a section close to that in *D* showing Golgi cisternae (green arrowheads) among immature mucin granules. (*F*) An image from a section adjacent to that in *E* showing Golgi cisternae (green arrowheads), with one of these adjacent to an immature mucin granule (red asterisk). (*G*) Reconstruction from multiple sections of a portion of the Golgi apparatus of a mouse secretory cell superimposed on the image shown in *F*. The nucleus is shown in blue, Golgi elements in green, and the immature secretory granule in red. (*H*) Reconstruction of different sections from those in *G* of the nucleus and Golgi apparatus superimposed on the same image. (*I*) The reconstructed nucleus and Golgi apparatus shown in isolation and rotated. It appears that the Golgi ribbon and nearby satellites may be connected, though the segmentation is not at sufficient resolution to be certain. (*J*) A high resolution image corresponding to Video 2 at 11 seconds showing four secretory cells containing mucin granules and one ciliated cell with a prominent ciliary tuft. Mature mucin granules near the cell apices contain electron-dense spherical regions, usually in the center of the granule (red arrowhead). Immature mucin granules are relatively electron-lucent (blue arrow), but maturing granules can show intermediate electron density and electron-dense central regions (red arrow). There is extensive endoplasmic reticulum (blue arrowhead) among the immature mucin granules One of the secretory cells, outlined with a dashed yellow line, appears to be undergoing exocytosis, with swelling of the central spherical region. (*K-L*) Images from sections at 13 and 18 seconds with the same annotation scheme as in *J*.

**Figure E6.**
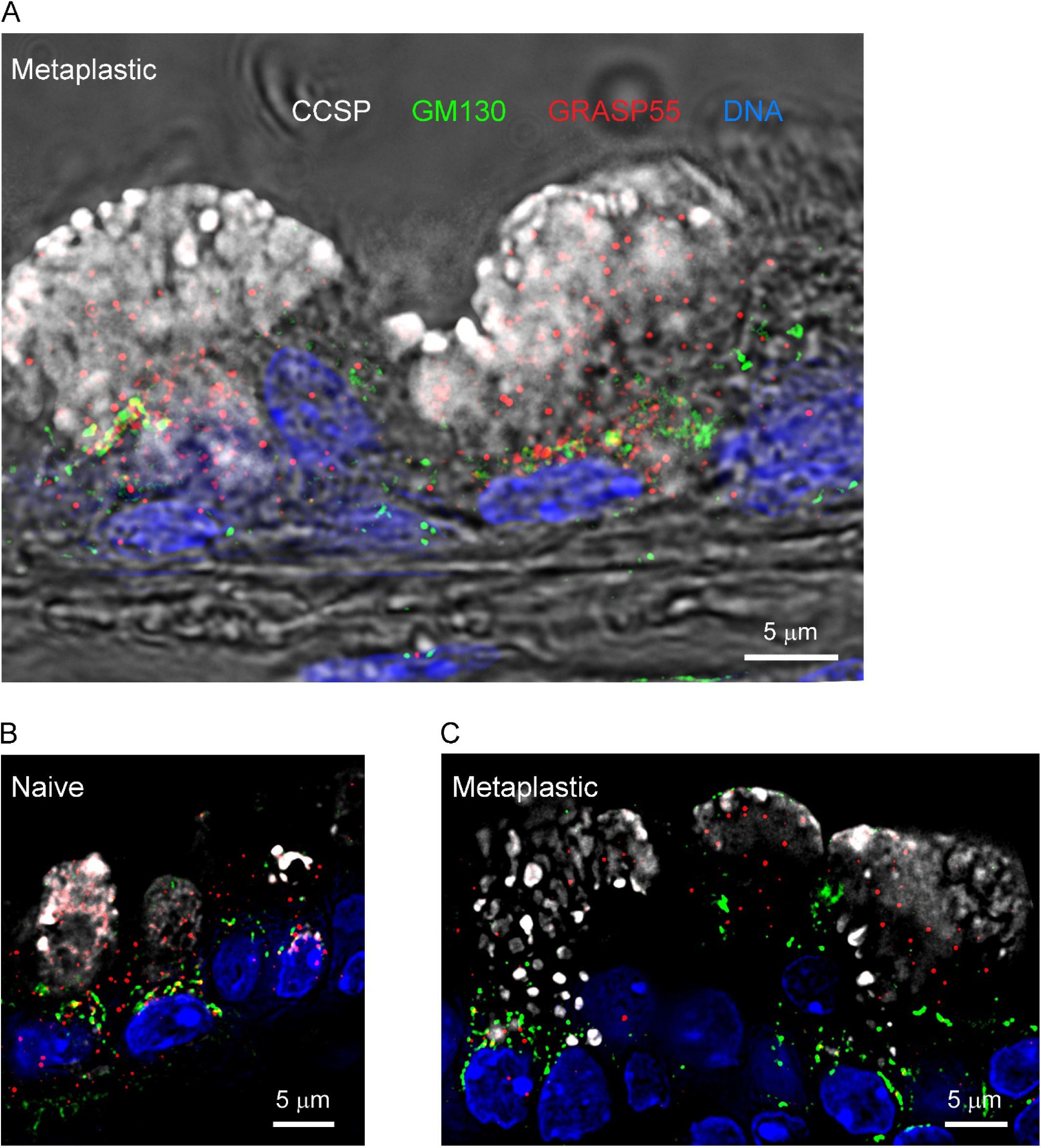
Additional images of the dissociation between cis and trans Golgi cisternae in mouse airway secretory cells. *(A)* Immunostaining and microscopy are exactly as in Figure 3 using antibodies against CCSP to mark secretory cells, GM130 to mark cis-Golgi cisternae, GRASP55 to mark trans-Golgi cisternae, and DAPI to mark nuclei in the axial bronchus of a mouse with mucous metaplasia due to treatment with IL-13.Fluorescence image is superimposed on a differential interference contrast (DIC) image. *(B)* As in *A*, but in a naïve mouse without mucous metaplasia, and without DIC image. *(C)* As in *A*, in another mouse with mucous metaplasia, without DIC image.

**Figure E7.**
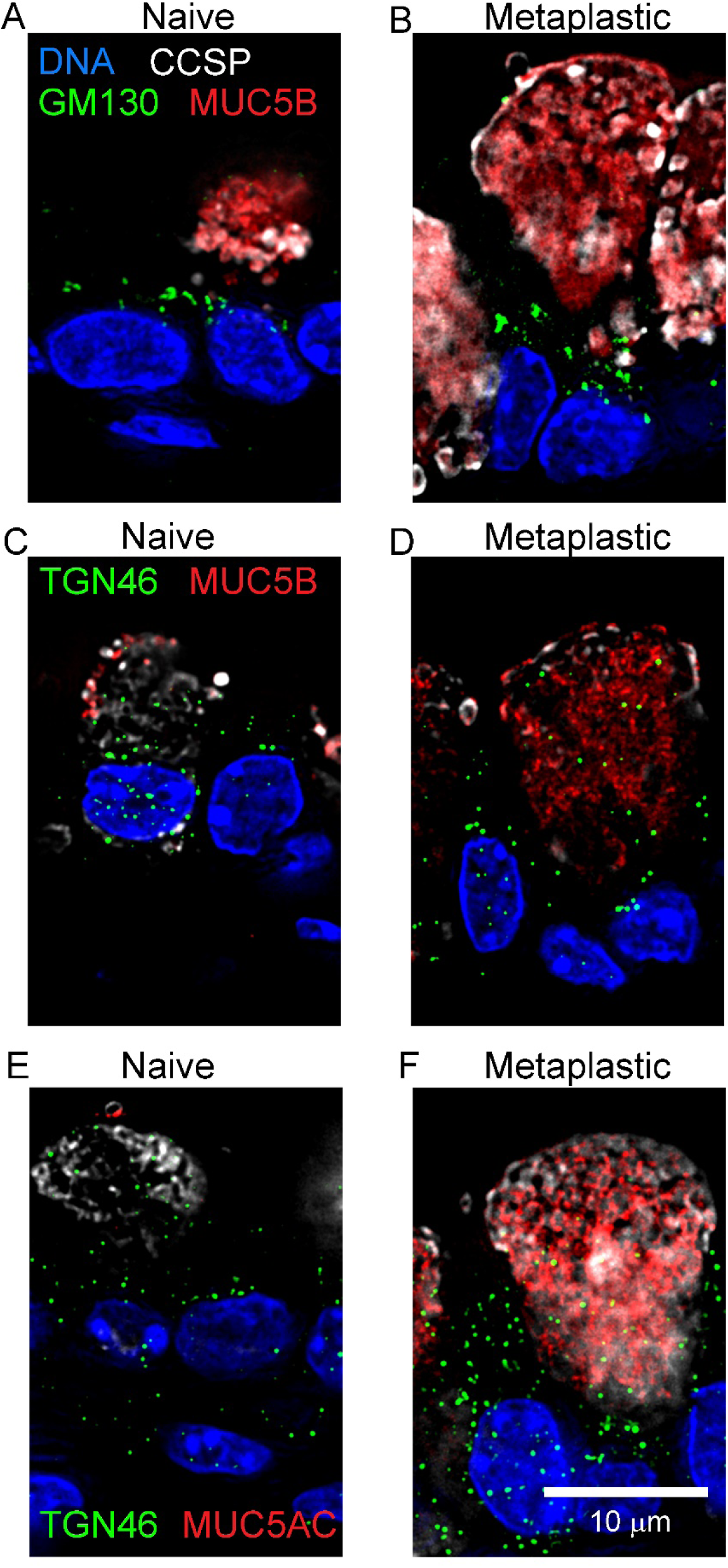
Localization of cis-Golgi cisternae and TGN with mucin granules in mouse airway secretory cells. *(A)* Immunofluorescence deconvolution microscopy using antibodies against CCSP to mark secretory cells, GM130 to mark cis-Golgi cisternae, MUC5B to mark mucin granules, and DAPI to mark nuclei in the axial bronchus of a naïve mouse airway. *(B)* Microscopy as in *A*, but in an airway with mucous metaplasia due to treatment with IL-13. *(C-D)* Microscopy as in *A-B*, but using TGN46 rather than GM130. *(E-F)* Microscopy as in *C-D* but using antibodies against MUC5AC rather than MUC5B.

**Figure E8.**
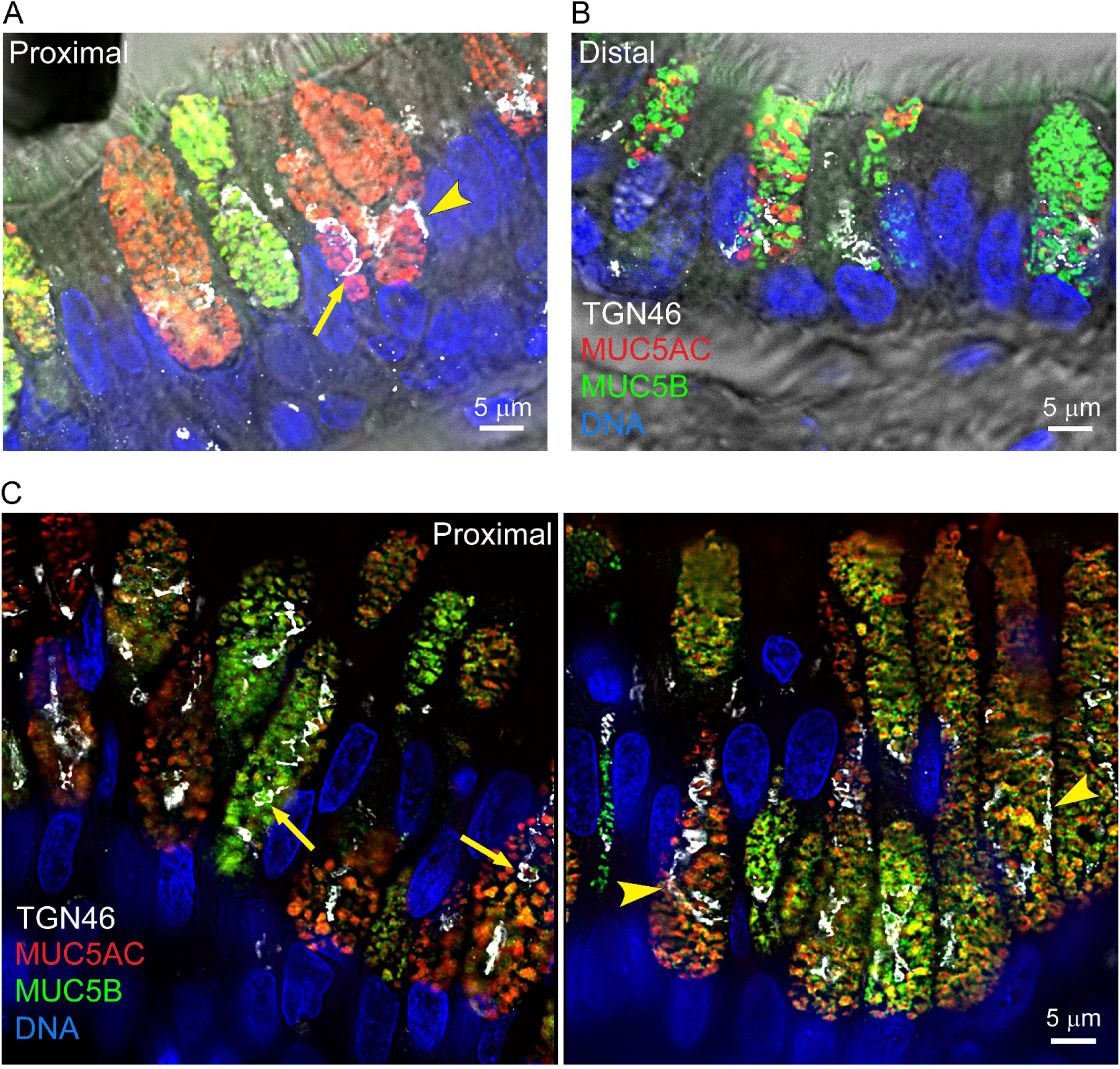
Additional immunofluorescence images of Golgi elements and mucins in human airway secretory cells. *(A)* Immunostaining and fluorescence microscopy as in Figure 4, but using antibodies against MUC5AC and MUC5B, together with antibodies against TGN46 to mark the trans-Golgi network and DAPI to mark nuclei in a proximal human airway. The fluorescent image is merged with a differential interference contrast (DIC) image. *(B)* Microscopy as in *A*, but in a distal human airway. *(C)* Two additional images of proximal human airways as in *A*, but without DIC images.

**Figure E9.**
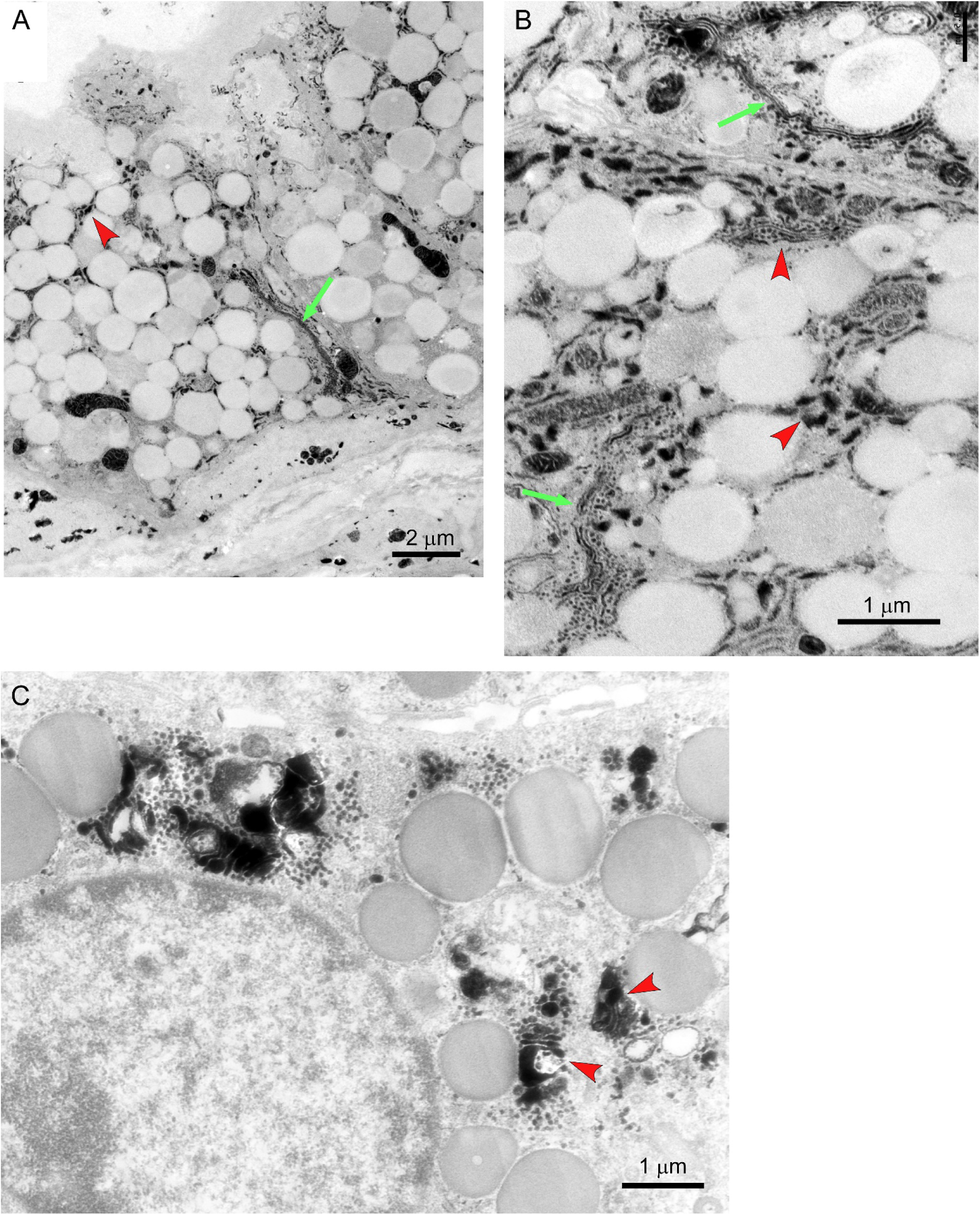
Additional EM images of Golgi ribbons and satellites in human submucosal gland mucous cells. Human airway tissue was fixed and stained with ZIO to highlight Golgi elements, as in Figure 5D. (*A*) A Golgi ribbon (green arrow) is seen extending from the nucleus alongside mucin granules, and Golgi satellites (red arrowhead) are seen among the granules. (*B-C*) Additional Golgi ribbons and satellites are indicated as in *A*.

**Figure E10.**
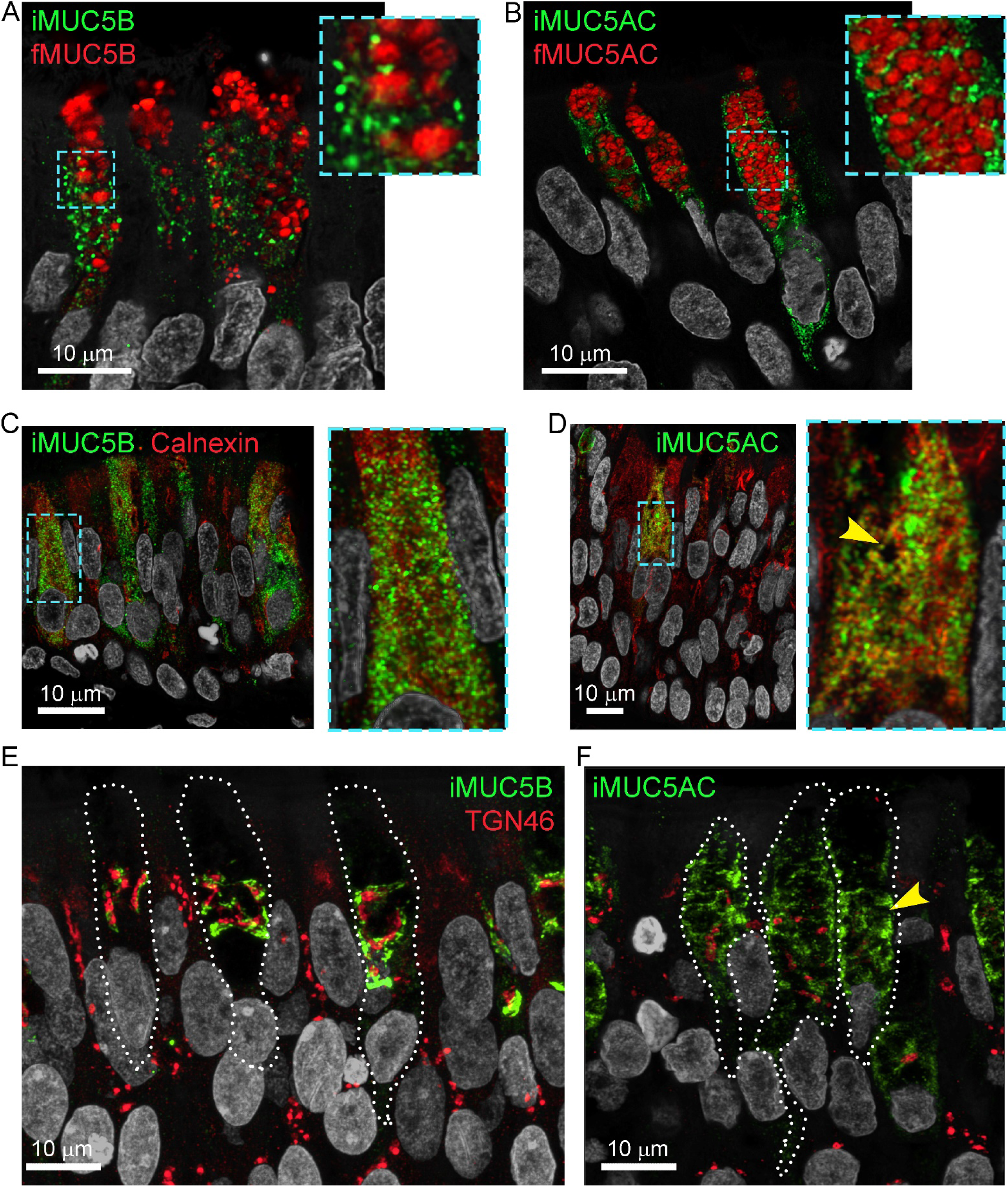
Distribution of incompletely and fully glycosylated mucins in human airway secretory cells. (*A-B)* Immunofluorescence Airyscan microscopy of normal human tracheal tissue using antibodies against incompletely glycosylated mucin proteins (iMUC5AC and iMUC5B), and against fully glycosylated and folded mucin proteins (fMUC5AC and fMUC5B) to mark granules. Hoechst stain (grey) marks nuclei. *(C-D)* Immunofluorescence microscopy as in *A-B*, but with antibodies against calnexin to mark ER instead of antibodies against fully glycosylated mucins. Yellow arrowhead points to black circle occupied by mucin granule. *(E-F)* Immunofluorescence microscopy as in *A-B*, but with antibodies against TGN46 to mark the trans-Golgi network instead of antibodies against fully glycosylated mucins. Yellow arrowhead points to black circle occupied by mucin granule.

**Figure E11.**
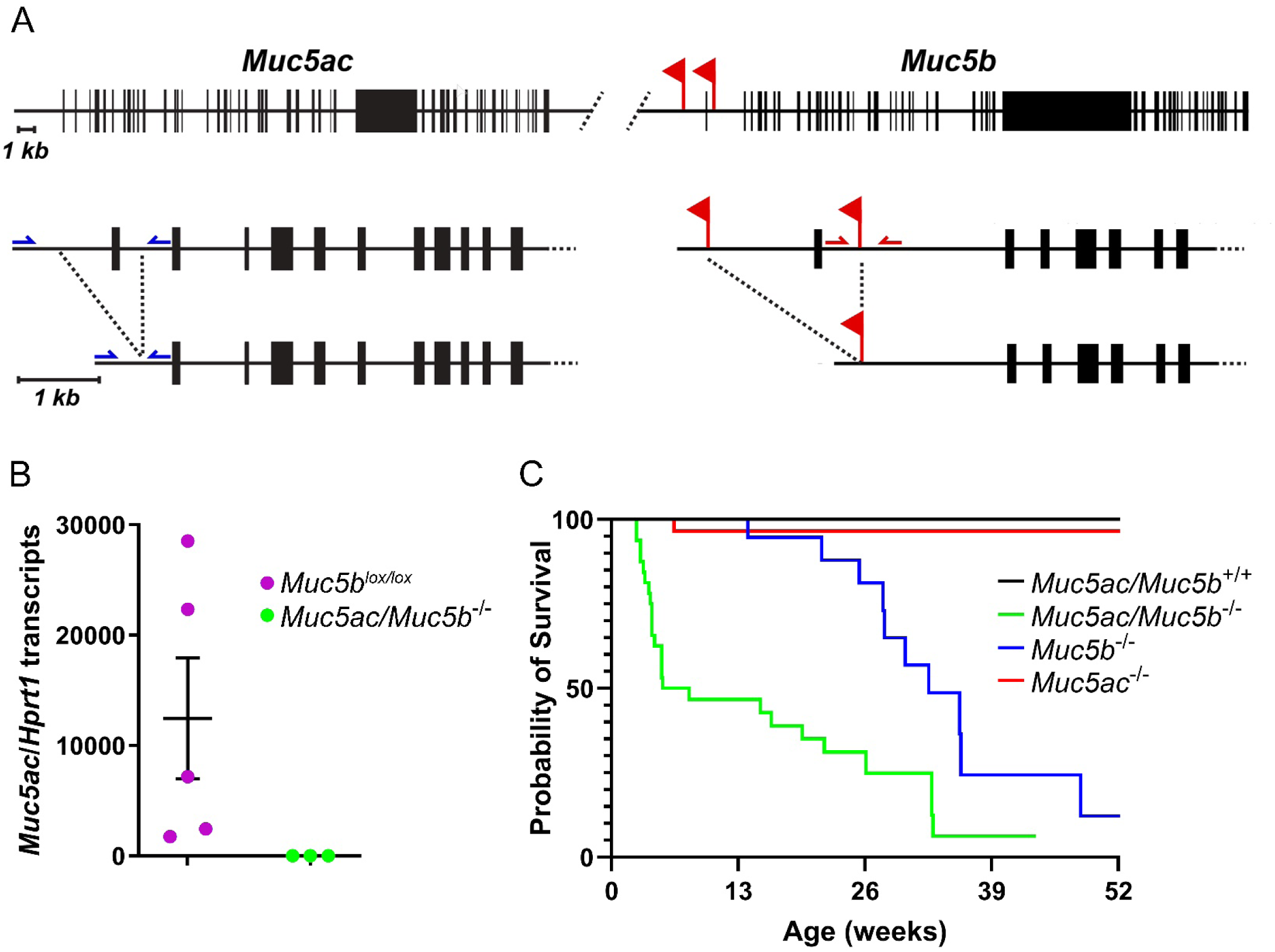
Generation of *Muc5ac/Muc5b* double deletant mice. *(A)* The *Muc5b* gene (right panel) was previously targeted to insert *loxP* sites (red flags) flanking the first exon. The gene structure is shown at low resolution in the top figure, at higher resolution in the middle figure, and after Cre-mediated recombination in the bottom figure. Mice bearing the floxed *Muc5b* allele were used for CRISPR/NHEJ-mediated excision of the first exon of *Muc5ac* (left panel). The location of primers used for genotyping is indicated by single-sided arrows. *(B) Muc5ac* transcripts relative to transcripts of the *Hprt1* housekeeping gene in the stomach of double deletant mice were compared to those in *Muc5b^lox/lox^* mice in which the *Muc5ac* gene was intact. (*C*) Post-natal survival is plotted of mice with intact or deletant alleles of *Muc5ac* and *Muc5b*.

**Video 1. SBF-SEM and reconstruction of the Golgi apparatus of a mouse secretory cell.** See Figure E5A-F for explanation. Best viewed with QuickTime. Not available in preprint.

**Video 2. SBF-SEM of multiple mouse secretory cells demonstrating ultrastructure of mucin granules.** See Figure E5J-L for explanation. Best viewed with Windows Player. Not available in preprint.

---

**Table S1.** Sources of human airway tissue at the University of Texas.

| **Table S1. Sources of human airway tissue at the University of Texas.** |  |  |  |  |  |  |  |  |
| --- | --- | --- | --- | --- | --- | --- | --- | --- |
| Study # | Age | Sex | Height | Weight | Lung | Lung dx | Smoking | Cause of death |
| 1065 | 35 | M | 178 | 95 | R | None | Unknown | Brain anoxia 2° drug intoxication |
| 1066 | 21 | F | 177 | 67 | R | None | Yes | Head trauma |
| 1069 | 20 | M | 180 | 112 | R | None | Unknown | Head trauma |
| 1071 | 29 | M | 170 | 70 | L | None | No | Brain anoxia 2° drug intoxication |
| **Height** is in cm; **Weight** is in kg. <u>Abbreviations:</u> **Lung dx** = known lung disease; **R** = right; **L** = left. |  |  |  |  |  |  |  |  |

**Table S2.** Antibodies used for immunofluorescence microscopy.

| **Table S2. Antibodies used for immunofluorescence microscopy.** |  |  |
| --- | --- | --- |
| **Antibodies** | **Source** | **Identifier** |
| Primary antibodies |  |  |
| Mouse monoclonal anti-MUC5AC | ThermoFisher | 45M1, # MA5-12178 |
| Mouse monoclonal anti-MUC5AC DyL550 conjugate | Novus Biologicals | 45M1, # NBP2-32732R |
| Rabbit polyclonal anti-MUC5B | Ehre laboratory | UNC222 (4) |
| Rabbit polyclonal anti-human MUC5B | MilliporeSigma | # HPA008246 |
| Rabbit polyclonal anti-human MUC5B | Hansson laboratory | MUC5B-D3 (5) |
| Mouse monoclonal anti-human iMUC5AC* | Clausen laboratory | CLH2 (6) |
| Mouse monoclonal anti-human iMUC5B<sup>†</sup> | Clausen laboratory | PANH2 (7) |
| Mouse monoclonal anti-AcTubulin | Sigma-Aldrich | T6793 |
| Goat polyclonal anti-uteroglobin | EMD Millipore | ABS1673 |
| Rat monoclonal anti-human uteroglobin | R&D Systems | MAB4218 |
| Goat polyclonal anti-mouse peptidase inhibitor 16 | R&D Systems | AF4929 |
| Rabbit polyclonal anti-MEOX2 | Novus Biologicals | NBP2-30647 |
| Rabbit polyclonal anti-mouse TGN46 | Abcam | ab16059 |
| Sheep polyclonal anti-human TGN46 | Bio-Rad | AHP500GT |
| Rabbit monoclonal anti-TGN46 | ThermoFisher | JF1-024, # MA5-32532 |
| Rabbit polyclonal anti-GRASP55 | Novus Biologicals | NBP1-89747 |
| Mouse monoclonal anti-GM130 | BD Biosciences | 610822 |
| Rabbit polyclonal anti-calnexin | Abcam | ab22595 |
| **Secondary antibodies** |  |  |
| Donkey anti-mouse, Alexa 488 | ThermoFisher | # A-21202 |
| Donkey anti-mouse, Alexa 555 | ThermoFisher | # A-31570 |
| Donkey anti-mouse, Alexa 647 | ThermoFisher | # A-31571 |
| Donkey anti-rabbit, Alexa 488 | Jackson Immuno | # 715-545-152 |
| Donkey anti-rabbit, Cy3 | Jackson Immuno | # 711-165-152 |
| Donkey anti-rabbit, Alexa 647 | ThermoFisher | # A-31573 |
| Donkey anti-rabbit, Alexa 555 | ThermoFisher | # A-31572 |
| Donkey anti-goat, Alexa 647 | Jackson Immuno | # 705-605-147 |
| Rabbit anti-sheep, unconjugated | Jackson Immuno | # 313-001-003 |
| If antibody is known to be species-specific towards its target, this is indicated.<br>*iMUC5AC = incompletely glycosylated MUC5AC, <sup>†</sup>iMUC5B= incompletely glycosylated MUC5B. |  |  |

